# Spike Protein Targeting “Nano-Glue” that Captures and Promotes SARS-CoV-2 Elimination

**DOI:** 10.1101/2021.04.13.439641

**Authors:** Guofang Zhang, Yalin Cong, Guoli Cao, Liang Li, Peng Yu, Qingle Song, Ke Liu, Jing Qu, Jing Wang, Wei Xu, Shumin Liao, Yunping Fan, Yufeng Li, Guocheng Wang, Lijing Fang, Yanzhong Chang, Yuliang Zhao, Diana Boraschi, Hongchang Li, Chunying Chen, Liming Wang, Yang Li

## Abstract

The global emergency caused by the SARS-CoV-2 pandemics can only be solved with adequate preventive and therapeutic strategies, both currently missing. The electropositive Receptor Binding Domain (RBD) of SARS-CoV-2 spike protein with abundant β-sheet structure serves as target for COVID-19 therapeutic drug design. Here, we discovered that ultrathin 2D CuInP_2_S_6_ (CIPS) nanosheets as a new agent against SARS-CoV-2 infection, which also able to promote viral host elimination. CIPS exhibits extremely high and selective binding capacity with the RBD of SARS-CoV-2 spike protein, with consequent inhibition of virus entry and infection in ACE2-bearing cells and human airway epithelial organoids. CIPS displays nano-viscous properties in selectively binding with spike protein (*K_D_* < 1 pM) with negligible toxicity *in vitro* and *in vivo*. Further, the CIPS-bound SARS-CoV-2 was quickly phagocytosed and eliminated by macrophages, suggesting CIPS could be successfully used to capture and facilitate the virus host elimination with possibility of triggering anti-viral immunization. Thus, we propose CIPS as a promising nanodrug for future safe and effective anti-SARS-CoV-2 therapy, as well as for use as disinfection agent and surface coating material to constrain the SARS-CoV-2 spreading.

## Introduction

The Corona Virus Disease 2019 (COVID-19), induced by severe acute respiratory syndrome coronavirus 2 (SARS-CoV-2), broke out since December 2019 and has become a worldwide health crisis^1^. The US FDA granted Remdesivir and hydroxychloroquine with Emergency Use Authorization to treat COVID-19, based on earlier reports of their capacity to inhibit SARS-CoV-2^2–4^, but revoked the use of hydroxychloroquine for COVID-19^5,6^, due to lacking effectiveness in clinical use^7,8^. Although Remdesivir has been approved for COVID-19 treatment in October 22^nd^, 2020, by US FDA, the interim results of the WHO Solidarity trial showed no significant effect^9–11^. Several mutations of SARS-CoV-2 on Spike (S) protein have been recently identified, which apparently cause increased viral infectivity^12^. The mutated SARS-CoV-2 variants have raised concerns regarding the effectiveness of the currently approved anti-SARS-CoV-2 drugs, neutralizing antibodies and vaccines^13^. Further, the resource-consuming logistics of cold-chain products risks to fail stopping SARS-CoV-2 transmission and pandemic spread^14^. Thus, despite a powerful effort in drug repurposing for the identification of new antiviral compounds^15,16^, novel compounds with effective antiviral activity are an urgent need for the treatment and containment of the SARS-CoV-2 pandemic infection^17^.

The host infection by SARS-CoV-2 requires the recognition and binding of the Receptor Binding Domain (RBD) of Spike (S) protein of SARS-CoV-2 to the host cellular ACE (angiotensin converting enzyme) 2 receptor^18,19^. S protein and its RBD thus serve as efficient targets for antiviral drugs. Anti-SARS-CoV-2 neutralizing monoclonal antibodies (nAbs) have been designed for targeted interaction with the RBD of S protein^20,21^ or the epitopes of RBD/ACE2 binding site^22–24^, or selected for potent neutralization activity^25–27^. However, the massive dosage of nAb for effective therapy (10 to 100 mg/kg), their uncertain effectiveness for mutated SARS-CoV-2 variants, and the complex nAb storage and shipping conditions necessary for their long-term integrity and efficacy, push to searching for alternative tools with a comparable therapeutic capacity^28^. In this view, the antimicrobial and antiviral capacity of nanomaterials (NMs)^29,30^ has raised particular interest in view of novel anti-SARS-CoV-2 strategies. NMs exhibit good capability in antiviral applications due to large surface area, tunable surface properties, and chemical reactivity. VivaGel, the best known antiviral dendrimer gel, has been used for HIV and HSV prevention by blocking the virus-cells interaction^31^. Metal NMs display antiviral capacity due to the interaction of metallic atoms with virus components, as in the case of silver NM^32^ and gold NM^33^. Gold NM with a tunable surface chemistry design that mimics heparan sulfate proteoglycan, effectively binding to viral attachment molecules, has been successfully applied to inhibit the infection of various viruses^30^. Thus, we propose to develop a S protein-targeting viscous nanodrug that could combine high anti-viral efficacy with excellent biocompatibility, easy preparation and convenient storage characteristics, as a promising new tool to prevent/treat COVID-19.

According to the Adaptive Poisson-Boltzmann Solver (APBS) electrostatics calculations^34^ and the crystal structure of the RBD and ACE2 interface^35,36^, the surface for RBD and is composed of abundant β-sheet structure, which serves as a target for efficient drug design. Considering structural similarity of β-sheet^37^, RBD preferentially adsorbs on the two-dimensional (2D) NM surfaces with abundant negative charges, which allowed for screening a series of NMs for antiviral activity (fig. S1-S3). We have discovered the copper indium thiophosphate (CuInP_2_S_6_, CIPS) nanosheets (NS), an ultrathin 2D and ferro-electric nanomaterial, which exhibits powerful anti-SARS-CoV-2 capacity. The CIPS NS can firmly adsorb the S protein RBD that duo to its physical nature including chemical component (Cu and S) and electronic structure (with a plenty of sulfur atoms, high average surface atomic and charge density, as well as the easy drifting of electrons for copper atoms in the crystal), and further induce RBD conformational change, successfully leading to the inhibition of SARS-CoV-2 infection of host cells. Macrophages have the capacity to eliminate NMs to protect the body integrity^38^. Thus, the SARS-CoV-2 trapped by CIPS NS was found to be largely phagocytosed by macrophages, thereby promoting viral elimination. Importantly, CIPS remains high affinity with mutant N501Y RBD of S protein, suggesting a consistent capacity on anti-SARS-CoV-2.

Its efficacy and excellent biocompatibility suggest CIPS NS as a promising anti-SARS-CoV-2 drug. The viscous flypaper-like and selective binding capacity of CIPS for the SARS-CoV-2 S protein also makes it particularly promising as surface coating material and disinfection agents to contain the SARS-CoV-2 spreading.

### The design and characteristic of CIPS

CIPS NS was exfoliated from the bulk CIPS (fig. S4a) with Li-intercalation by using n-butyl lithium and ultrasound sonication, and its crystalline nature was proven by X-ray diffraction (XRD) characterization (fig. S4b). The exfoliated CIPS showed a multi-layer structure with a mean size of ~200 nm determined by SEM (fig. S4c) and TEM (Fig. 1a), and an average thickness of ~3.1 nm by AFM (Fig. 1b–c). According to extended X-ray absorption fine structure (EXAFS), copper atoms coordinate with sulfur atoms (Fig. 1d, fig. S4d-g and Table S1), where Cu has a valence of +1 (Table S2). Sulfur was present as sulfide (S^2-^) with a valence of −2 (fig. S4h-i). According to In L3-edge XANES, Indium mainly forms In-S, In-P, and In-O bonds, and exhibits +3 valence (fig. S4j). The chemical form of In is consistent with crystal structure of CIPS NS where In atoms remain between two different CIPS surface and stably support the structure of CIPS by coordinating with the P and S atoms (Fig. 1e). The structure of CIPS NS is described in the schematic diagrams in Fig. 1e, which show the Cu, S, and P atoms mainly localized on the surface and the In atoms immersed in the NS.

**Fig. 1.**
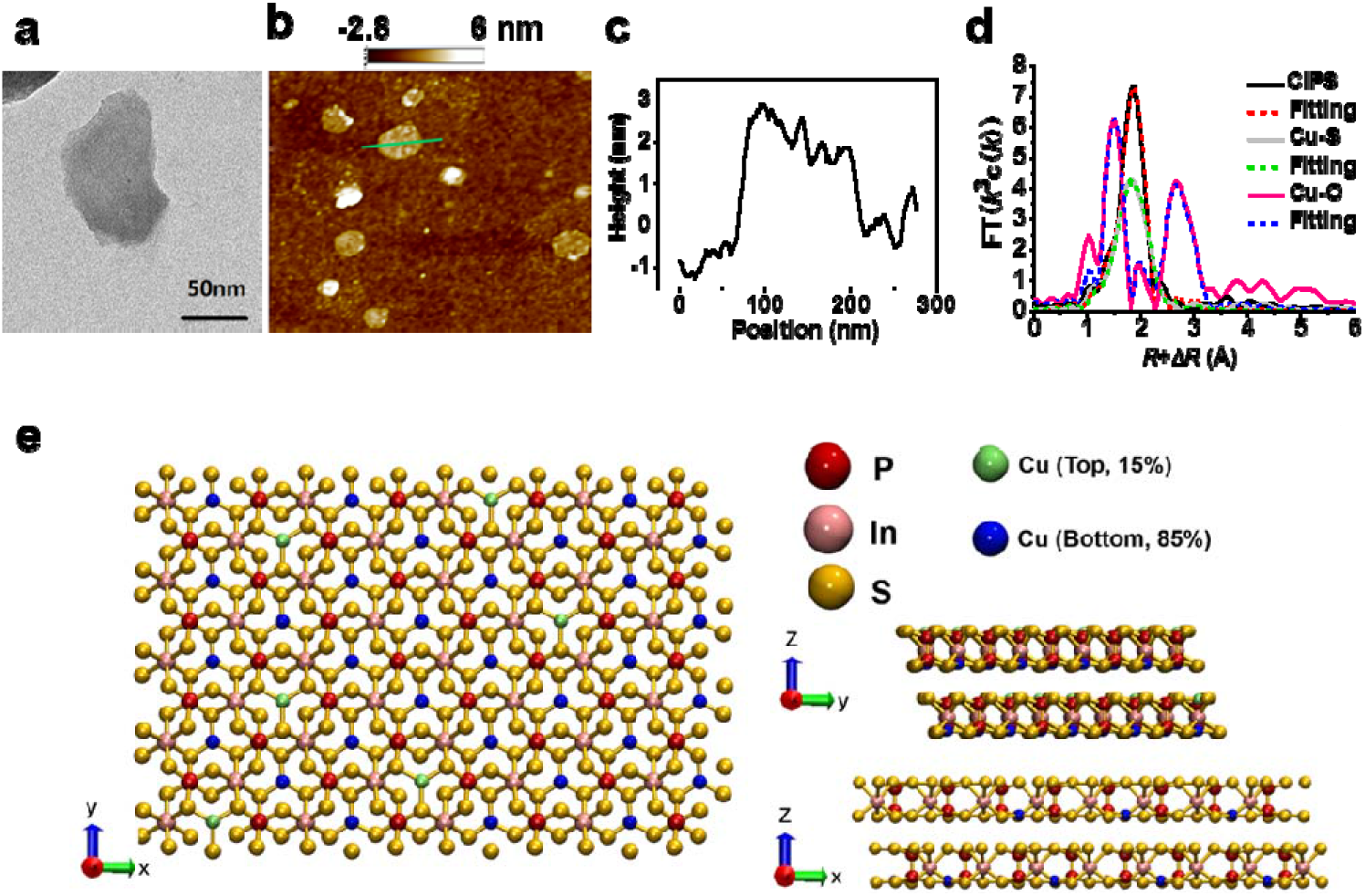
Characterization of CIPS NS. a) Representative TEM image of exfoliated CIPS NS. b-c) AFM image of CIPS NS showing thickness and size distribution. d) Coordination structure of Cu with S atoms in CIPS as determined by EXAFS. e) Schematic illustration of the CIPS crystal structure from top, left and right sides.

### CIPS efficiently inhibits the infection of SARS-CoV-2 virus

The anti-SARS-CoV-2 effect of CIPS NS (shorter as CIPS) was investigated with the established pseudoviruses SC2-P (expressing the SARS-CoV-2 S protein) and SARS-P (expressing the SARS S protein). CIPS showed an excellent biocompatibility with several human/monkey epithelial and myeloid cell lines (fig. S5). The capacity of CIPS to inhibit SC2-P infection was dose-dependent, as shown with a luciferase reporter assay in both Vero-E6 and ACE2/293T cells (Fig. 2a) and in immunofluorescence (IF) in ACE2/293T (Fig. 2b; quantitative evaluation in Fig. 2c). The CIPS inhibitory effect was evident at early time points (0.5-2 h) (Fig. 2d–e) and persisted up to 15 h (fig. S6). Although we could detect many SC2-P particles around cells (fig. S6a-c), the virus apparently cannot enter and infect cells, as indicated by luciferase activity (fig. S6d).

**Fig. 2.**
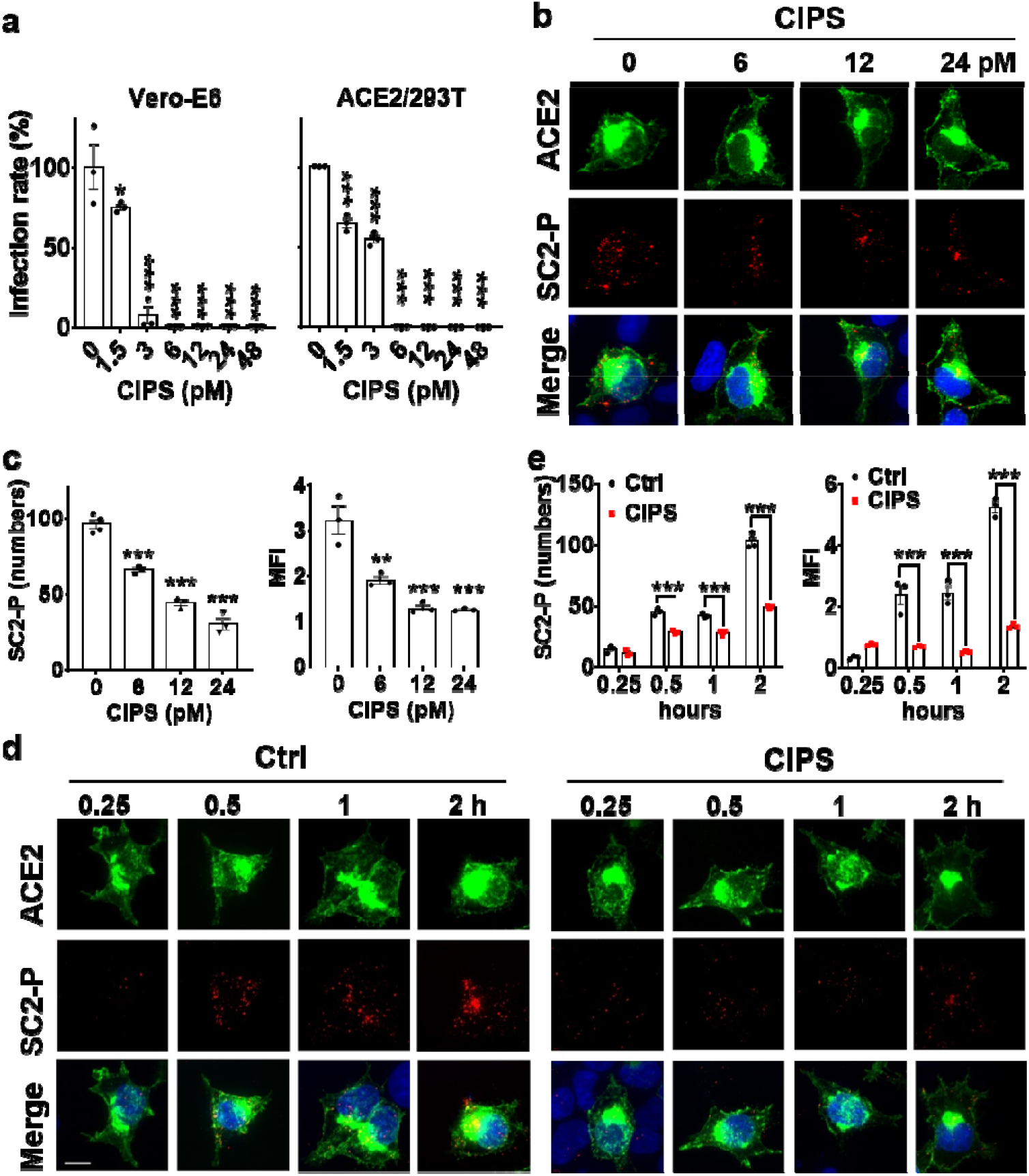
CIPS inhibits the SC2-P infection. a) The inhibitory effect of CIPS on SC2-P infection in Vero-E6 and ACE2/293T cells. SC2-P was pre-incubated with 0-48 pM CIPS for 2 h and added to Vero-E6 or ACE2/293T cells for 2 h, then washed off. Infection was evaluated after 40 h on cell lysates as SC2-P dependent luciferase activity. Data are representative of three independent experiments, and presented as mean ± SEM of technical triplicates. *, p<0.05; ***, p<0.005 by ANOVA. b-e) Dose-dependency (b-c) and time course (d-e) of the inhibitory effect of CIPS on SC2-P infection of ACE2-OE cells. SC2-P was pre-incubated with CIPS for 2 h and added to ACE2-OE cells for 2 h (b); or pre-incubated with 12 pM CIPS for 2 h, and added to ACE2-OE cells for different times (d). Green: ACE2-GFP; red: SC2-P labeled with anti-Flag antibodies. c, e) Quantitative and statistical analysis of IF data. The number of SC2-P within infected ACE2-OE cells was counted (left, cells ≥ 3) and analyzed for mean fluorescent intensity (MFI, right). Data are from one out of three experiments performed, and presented as mean ± SEM of three representative cells. **, p<0.01; ***, p<0.005 by Student’s *t-*test.

We further assessed the anti-viral effects of CIPS with the authentic SARS-CoV-2 virus. After 48 h from SARS-CoV-2 infection of Vero-E6 cells, the SARS-CoV-2 infection rate was decreased by the presence of CIPS in a dose-dependent fashion (Fig. 3a–b and fig. S7), while cell viability was not affected, with a good selectivity index (SI; ratio between antiviral and cytotoxic concentrations) exceeding 17.6 (Fig. 3b). The phase contrast images in Fig. 3c also show a good anti-viral effect of CIPS, which protected the cells from the virus-induced cytopathic effect. To further evaluate the CIPS antiviral effect on infected cells, SARS-CoV-2 infected Vero-E6 cells were treated with CIPS, and the intracellular SARS-CoV-2 was effectively inhibited by CIPS in a dose dependent manner (Fig. 3d).

**Fig. 3.**
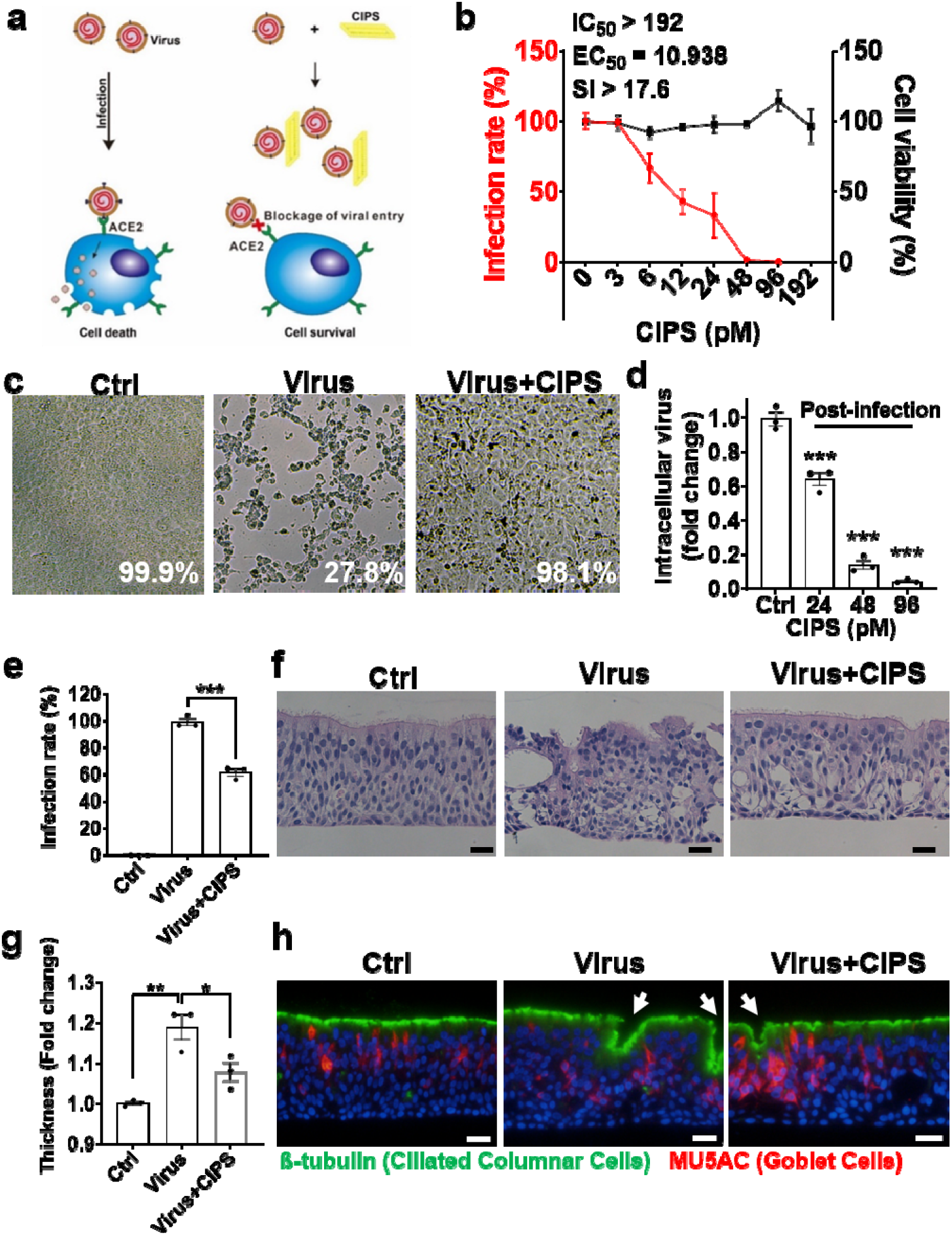
CIPS inhibits the SARS-CoV-2 infection. a) The schema of the CIPS blocking the infection of SARS-CoV-2 to the host cells. b) The antiviral effect of CIPS on SARS-CoV-2 infection with Vero-E6 cells was measured after 48 h virus post-infection with/without CIPS. Real-time PCR data for SARS-CoV-2 ORF1ab from one out of three experiments are presented as mean **±** SEM of 3-5 biological replicates. Cell viability of CIPS was evaluated after 48 h. c) Representative images of Vero-E6 cells 48 h post-infection with SARS-CoV-2. Ctrl: uninfected cells; Virus: cells infected with SARS-CoV-2; Virus + CIPS: cells infected with SARS-CoV-2 treated with 24 pM CIPS. d) Antiviral effect of CIPS on Vero-E6 cells with SARS-CoV-2 post-infection. Vero-E6 cells were firstly challenged with SARS-CoV-2 (12000 pfu) for 2 h, and continuously cultured in fresh or CIPS containing medium for another 48 h. Intracellular SARS-CoV-2 was evaluated by qPCR. e-h) The anti-SARS-CoV-2 capacity of CIPS on human airway epithelial organoids derived from human nasal tissue. (e) SARS-CoV-2 replication (ORF1ab) was observed upon infection on the cultured airway epithelium in the presence or absence of 24 pM CIPS. Data presented as mean ± SEM (n=3). ***, p<0.005 by Student’s *t-*test. (f) Representative image of H&E staining of cultured tissue cross-sections. Ctrl: uninfected tissue; Virus: tissue infected with SARS-CoV-2; Virus + CIPS: tissue infected with SARS-CoV-2 treated with 24 pM CIPS. (g) The thickness of the human respiratory epithelium section that measured by ImageJ. Data are representative of best epithelium section, and presented as mean±SEM of technical triplicates. *, p<0.05 and **, p<0.01 by Student’s t-test. (h) Cell distribution in the human respiratory epithelium after SARS-CoV-2 infection. Cross-section of the human respiratory epithelial tissue in control culture conditions (Ctrl), upon exposure to SARS-CoV-2 (Virus) and exposed to SARS-CoV-2 together with CIPS (Virus+CIPS). Green: tubulin staining the ciliated columnar epithelial cells; Red: MU5AC staining of mucus-producing goblet cells; Blue: DAPI staining of nuclei. Scale bar = 20 μm. The white arrows show the sites of tissue damage.

The anti-SARS-CoV-2 effect of CIPS was also assessed with air-liquid-interface (ALI)-cultured human airway epithelial organoids (Fig. 3e–h). Mature airway organoids presented a multi-layer airway epithelial structure with ciliated cells at the air-facing side. CIPS treatment showed a 40% decrease of SARS-CoV-2 replication (Fig. 3e), together with an effective protection on the tissue architecture and integrity, which on the contrary was severely damaged by SARS-CoV-2 infection (Fig. 3f–h). A thick airway epithelium can occur upon viral infection duo to an expanded intercellular space and reduced cell-cell adhesion, which caused by virus for their escape^39^. SARS-CoV-2 infection on human airway epithelial organoids caused an increased thickness of the epithelium, which can be reduced by CIPS treatment (Fig. 3g). Thus, these results indicate that CIPS displays an effective anti-SARS-CoV-2 capacity both on Vero-E6 cells and on *ex vivo* reconstructed human airway epithelial organoids in culture.

### The inhibition mechanism of CIPS on SARS-CoV-2 host infection

After having excluded possible metallic ion effects (fig. S8), we assessed the interaction of CIPS with SC2-P by TEM (Fig. 4a), and examined the variations in the CIPS physico-chemical properties (fig. S9). We further detected the interaction of CIPS with SC2-P by examining its S protein in Western blotting (Fig. 4b). Further, the interaction of CIPS with authentic SARS-CoV-2 virus was also analyzed, indicating that CIPS could effectively adsorb and trap SARS-CoV-2 virus (Fig. 4c). The capacity of CIPS to adsorb the S protein was also proven by Western blotting (Fig. S10). The analysis of CIPS binding to the RBD of the S protein by Biolayer Interferometry (BLI) shows a very high binding affinity (Fig. 4d, *K*_D_ <0.001 nM), in contrast with an at least 100x lower affinity for a number of serum proteins and factors (Fig. 4e and Table S3), suggesting a selective binding capacity for the S protein RBD. A mixture of SC2-P with several fold excess volume of FBS or BSA was used to evaluate the CIPS anti-viral effects in complex media, and shows that CIPS retains a strong binding capacity for the S protein of SC2-P (Fig. 4f), and that its anti-viral effect is not affected (Fig. 4g and fig. S11). The capacity of other 2D NMs (MoS_2_ and GO) to bind RBD (*K*_D_ 11.7 and 5.2 nM, respectively) was significantly lower than that of CIPS (fig. S12). The NM binding affinity for RBD (*K*_D_: CIPS≪GO< MoS_2_) is comparable to their anti-viral activity (fig. S3b). Binding of RBD to ACE2 (*K*_D_ 8.08 nM, in agreement with reported data^40^) was decreased almost 10 times if RBD was pre-exposed to CIPS, showing a *K*_D_ of 74.7 nM (Fig. 4h–i). All these data suggest that the selective and preferable binding to the S protein RBD makes CIPS a potentially effective neutralizing drug for COVID-19 treatment.

**Fig. 4.**
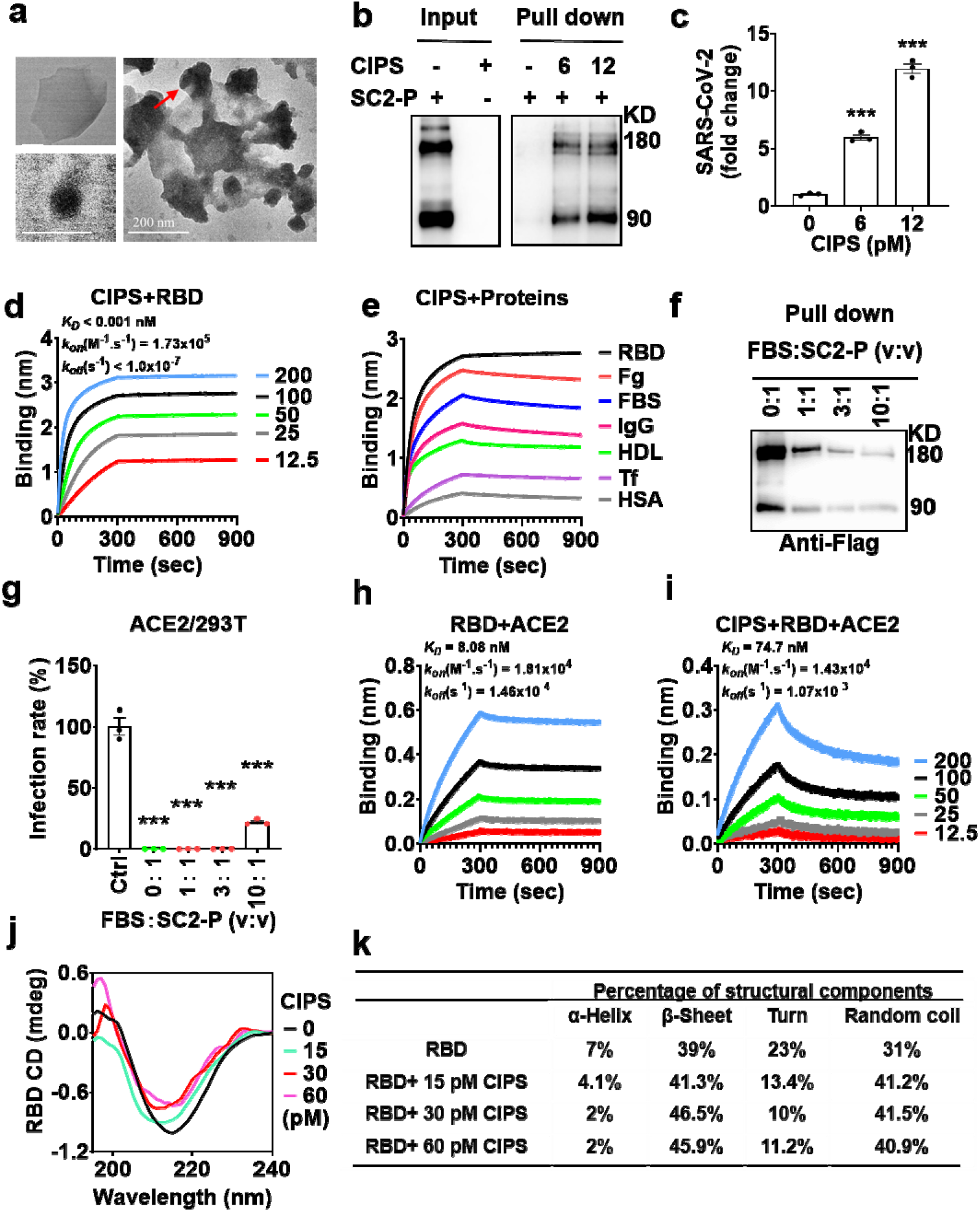
The SARS-CoV-2 inhibition mechanism of CIPS. a) TEM images of SC2-P, CIPS and SC2-P incubated with CIPS. b) WB analysis showing SC2-P binding to CIPS (pM). c) CIPS effectively binds to SARS-CoV-2 virus. CIPS incubated with SARS-CoV-2 for 2 h and separated by centrifugation. The SARS-CoV-2 was quantitatively assessed in CIPS precipitates. Data as mean ± SEM of technical triplicates. ***, p<0.005 by Student’s *t-*test. d, e) Binding affinity for CIPS of RBD at different concentrations (nM) (d) and various proteins at 50 nM (e) as determined by Biolayer Interferometry (BLI). f-g) The SARS-CoV-2 inhibiting capacity in complex biological matrices. SC2-P were mixed with FBS at different ratios and incubated with 12 pM CIPS for 2 h. f) WB analysis showing the specific binding of SC2-P to CIPS in the presence of FBS. g) Anti-SARS-CoV-2 activity of CIPS in the presence of FBS. The FBS/SC2-P/CIPS were used to infect ACE2/293T cells for 2 h. Cells were lysed after 40 h incubation and the intracellular SC2-P were detected based on luciferase activity. Data in (g) are from a representative experiment out of three performed, and presented as mean ± SEM of technical triplicates. ***, p<0.005 by Student’s *t-*test. h-i) Inhibition of RBD binding to ACE2 (nM, on the right) by CIPS, as determined by BLI. j-k) Conformational variations in the RBD structure after CIPS interaction, as determined by CD spectra.

A molecular dynamics (MD) simulation was performed to study the adsorption process of RBD on the CIPS surface. The crystal structures of ACE2 (cyan), S protein (brown), and the RBD (yellow) were used to elucidate their binding interface (fig. S13). The amino acid residues of RBD bound to ACE2 are listed in Table 1 and mostly locate in β-sheet, turn, and random coil stretches. The secondary structure of RBD in the presence and absence of 30 pM CIPS was characterized with circular dichroism (CD) spectra, and shows a majority of β-sheet conformation within the RBD structure (Fig. 4j–k). Since β-sheet structures promote protein adsorption on the NS^41^, this may explain the preferential binding of RBD to CIPS. A conformational change in RBD upon CIPS binding was also observed (Fig. 4k), with a decrease of α-helix (7% to 2%) and turn (23% to 10%), and an increase of β-sheet (39% to 46.5%) and random coil (31% to 41.5%). MD simulation was further used to explore binding interface of RBD to CIPS within 100 ns (fig. S14-S15, Table S4), and the typical configuration was shown (Fig. 5a), with the amino acid residues and the interactive forces contributing to the binding illustrated in colors (Fig. 5a). Among the 11 amino acid residues of RBD interacting with the ACE2, six showed strong adsorption with CIPS (Table 1, indicated by *).

**Table 1.** The binding residues between RBD and ACE2.

| Number | Residues in RBD | Residues in ACE2 | Conformational location in RBD | Interaction forces |
| --- | --- | --- | --- | --- |
| 1 | **LYS417\*** | ASP30 | Turn | Salt bridge |
| 2 | TYR449 | GLU37 | Coil | Hydrogen bond |
| 3 | TRY453 | HIS34 | β-Sheet | Hydrogen bond |
| 4 | LEU455 | HIS34 | Coil | Hydrophobic interaction |
| 5 | GLN474 | GLN24 | Coil | Hydrogen bond |
| 6 | **PHE486\*** | MET82 | Turn | Hydrophobic interaction |
| 7 | ASN487 | GLN24 | Turn | Hydrogen bond |
| 8 | **GLN493\*** | GLU35 | β-Sheet | Hydrogen bond |
| 9 | **GLN498\*** | GLN42 | Turn | Hydrogen bond |
| 10 | **THR500\*** | ARG357 | Turn | Hydrogen bond |
| 11 | **ASN501\*** | LYS353 | Turn | Hydrogen bond |

\* These residues consistently interact with CIPS.

**Fig. 5.**
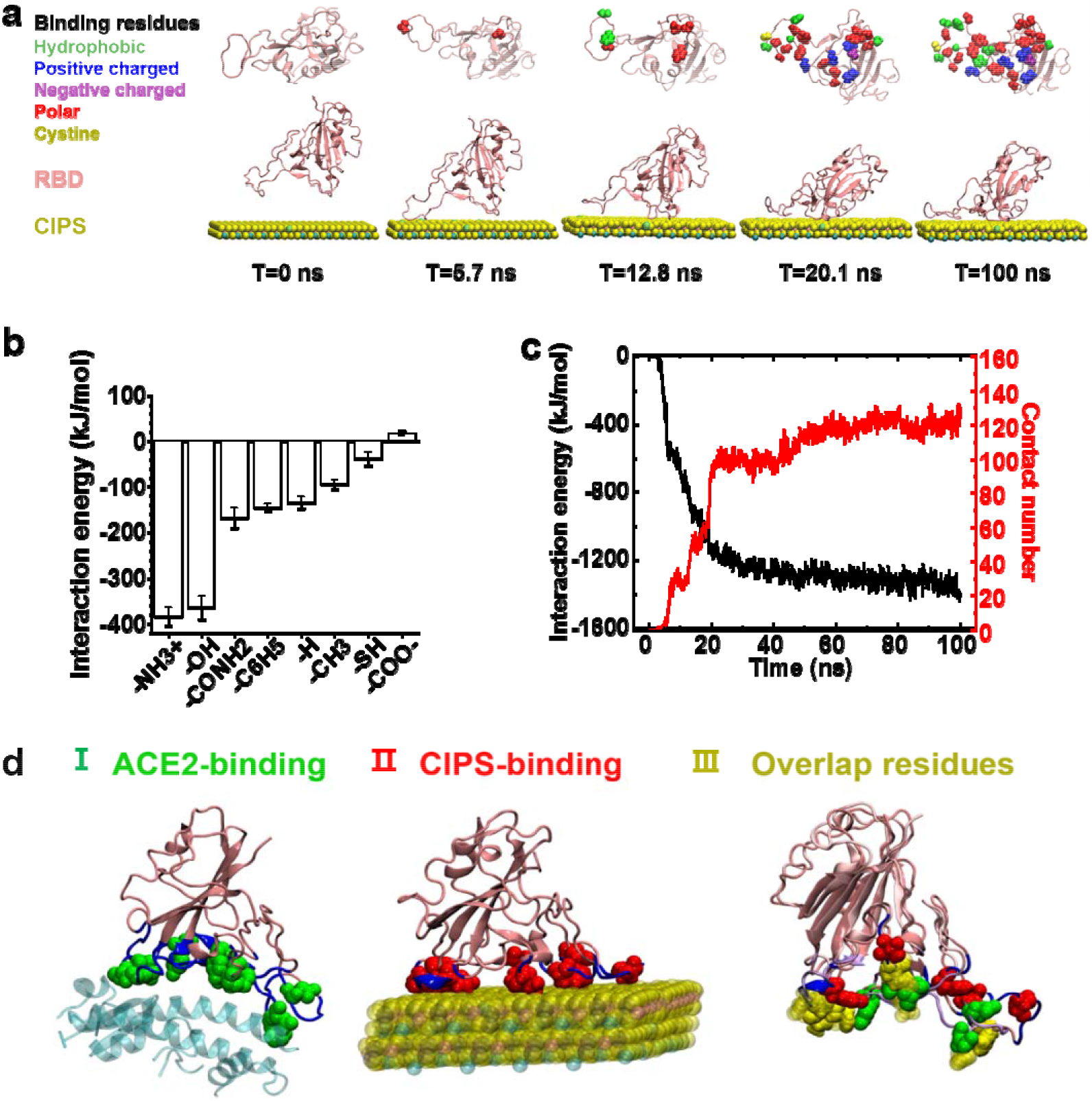
The simulation of CIPS binding to RBD. a) Snapshots from a representative trajectory for RBD and CIPS adsorption in System 1 based on MD simulation. The bottom row indicates the dynamic process of RBD adsorption on the surface of CIPS. The upper row shows the bottom view of the amino acid sites of RBD that bind with CIPS, where van der Waals spheres are the residues contacting CIPS (red, polar residues; green, hydrophobic residues; blue, positively charged residues; purple, negative residues; yellow, cysteines containing disulfide bonds). b) Preferred functional groups of RBD amino acids bound to the surface of CIPS, as determined by the interaction energies between CIPS and different functional groups based on MD simulations. c) The interaction energy between RBD and CIPS (black) and the number of contact atoms (with the exception of of H; red) in RBD when approaching the CIPS surface in System 1. d) CIPS binding to RBD interferes with RBD binding to ACE2. Amino acid residues at the RBD binding sites for ACE2 (a, green) and those of RBD to CIPS (at 100 ns in System 1; b, red) are largely overlapping (c, yellow).

Hydrophobic, polar and positively charged residues in RBD mainly contribute to the adsorption of RBD on CIPS (Table S5). The functional groups (−NH_3_^+^, −OH, −CONH_2_, −C_6_H_5_, −H, −CH_3_, and −SH) within amino acid resides take part in the adsorption of RBD to CIPS, whereby, based on the calculated interaction energy, the positively charged residues (Arg, Lys: −NH_3_^+^) and the polar ones (Ser, Tyr, Thr: −OH) could be considered as the most important interacting residues (Fig. 5b). The adsorption reaches equivalence after ~20 ns during the 100 ns simulation (Fig. 5c). Among the amino acid residues critically involved for the RBD/ACE2 binding (in green, Fig. 5d-(I)), six out of eleven are located in turn regions of RBD (Fig. 5d-(II), Table 1, Table S5) at 100 ns simulation (in red), depicted as the residues in yellow shared in both systems (Fig. 5d-(III)). Since interaction with CIPS decreases the turn conformation within RBD (evaluated by CD spectra; Fig. 4j–k), turn conformation will be probably disrupted, a circumstance that may change the binding interface of RBD with ACE2, thereby inhibiting the virus interaction with host cells and preventing infection.

The new variant of SARS-CoV-2, VOC-202012/01, possess a mutation of N501Y in the RBD domain that accounted for SARS-CoV-2 spreading^12^. MD simulation was also performed to analyze the interaction between CIPS and N501Y RBD. Although the local mutation residue from Asn to Tyr affects the binding energies of other residues with CIPS, the interaction energy of CPIS with N501Y RBD (−1218.06 ± 37.75 kJ/mol) is similar as that of wild type RBD (−1217.73 ± 49.97 kJ/mol) (fig. S16), suggesting that CIPS still maintains high affinity with the mutant N501Y RBD.

MD simulations also showed that, compared to MoS_2_ and GO NS binding to RBD (fig. S17-18 and Table S6), CIPS exhibits the lowest interaction energy (fig. S18b), the highest contact atom number (fig. S18c), and the largest contact area (fig. S18d). The average surface atomic density and the average surface charge density of CIPS are 12.863/nm^2^ and 6.262/nm^2^, respectively, much higher than those of MoS_2_ and GO. Thus, the highest binding affinity of CIPS to RBD can be also attributed to its strongest electrostatic attraction, with simulation data (fig. S18) being consistent with BLI results (fig. S12) and infection results (fig. S3).

### CIPS promotes phagocytosis and elimination of SARS-CoV-2 by macrophages

NMs can be eliminated by macrophages. Because of the strong interaction of CIPS with SARS-CoV-2, we investigated the possibility that macrophages could efficiently capture and eliminate SARS-CoV-2 when the virus is bound to CIPS. As shown in the figure 6 (a, b), CIPS-treated SC2-P could be effectively phagocytosed by macrophages (differentiated human THP-1 cells) after 24 h incubation, and the virus was effectively eliminated by macrophages during the subsequent degradation process (Fig.6a–b). SC2-P was found to be accumulated in macrophages after inhibiting the lysosome function with bafilomycin (BM) (Fig.6a–b), indicating the majority of SC2-P was degraded in lysosomes. The same findings were obtained with authentic SARS-CoV-2 virus (Fig. 6c–d). SARS-CoV-2 could be effectively eliminated after remove the extracellular virus (Fig. 6c). Lysosome inhibition with BM significantly increased the intracellular SARS-CoV-2 levels, implying the SARS-CoV-2 elimination was lysosome-dependent (Fig. 6d). Similar to the pseudovirus, SARS-CoV-2 phagocytosis was also greatly increased in the presence of CIPS (Fig. 6d).

**Fig. 6.**
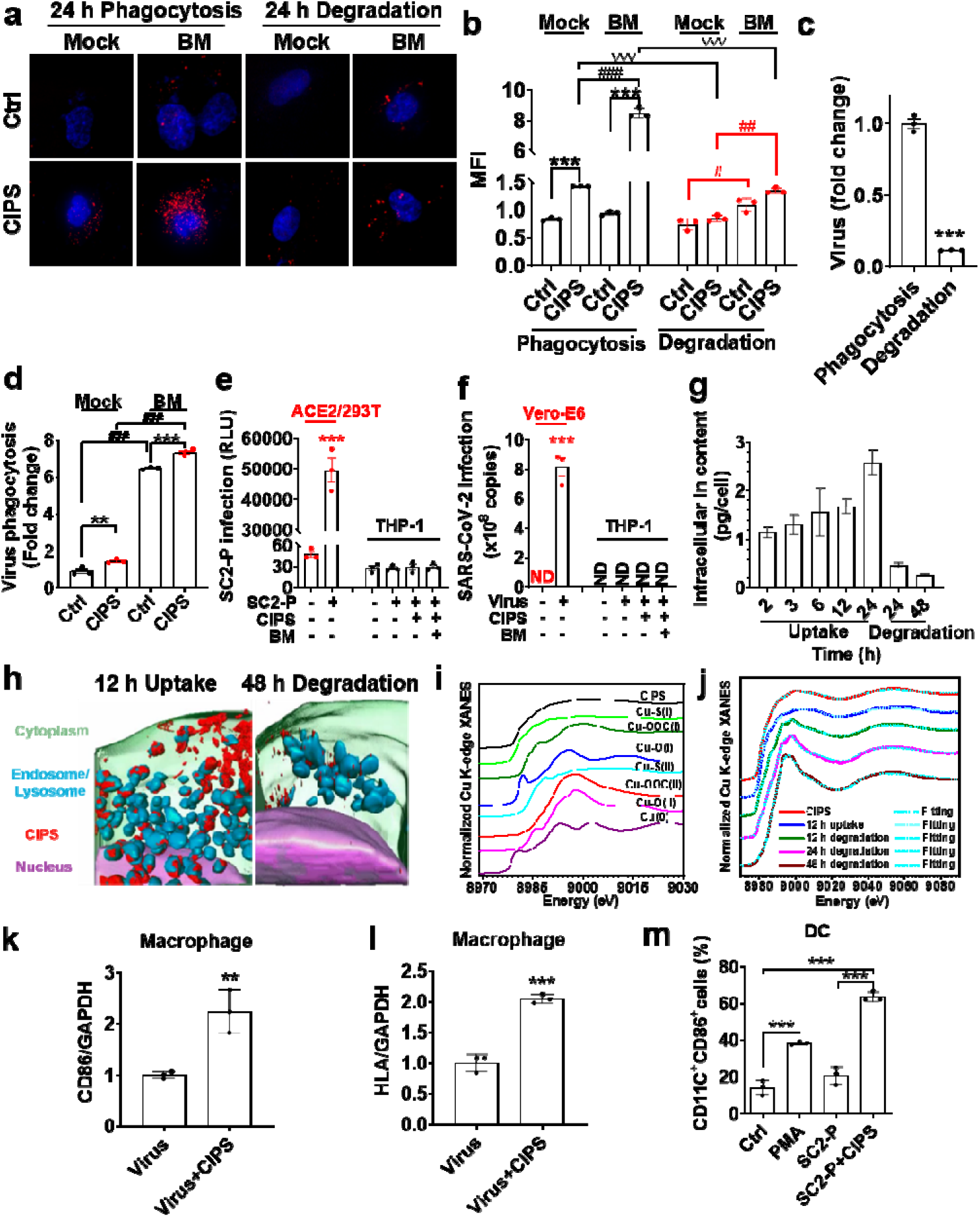
CIPS promoted the phagocytosis and elimination of SRAS-CoV-2 by macrophages. a-d) CIPS promotes the phagocytosis and elimination of SC2-P (a-b) and SARS-CoV-2 (c-d) in Macrophage-like differentiated THP-1 cells. a-b) The SC2-P were pre-incubated with 12 pM CIPS for 2 h. Macrophage-like differentiated THP-1 cells were treated with SC2-P with/without CIPS pre-incubation for 24 h (Phagocytosis). SC2-P containing medium was replaced by fresh medium and cultured for another 24 h (Degradation). b) Quantitative and statistical analysis with the infected SC2-P of IF data. c) Differentiated THP-1 was challenged with SARS-CoV-2 for 4 h (Phagocytosis). The SARS-CoV-2 challenged THP-1 was continuously cultured in fresh medium for another 48 h (Degradation). Intracellular SARS-CoV-2 was evaluated by qPCR. d) SARS-CoV-2 was pre-incubated with 12 pM CIPS for 2 h, and added to differentiated THP-1 cells for 48 h in the presence or absence of Bafilomycin (BM). Intracellular SARS-CoV-2 virus was assessed by qPCR. e-f) The SC2-P (e) and SARS-CoV-2 (f) cannot infect the differentiated THP-1 cells. Differentiated THP-1 was co-cultured with SC2-P (24 h) or SARS-CoV-2 (4 h), and the cultured mediums were replaced with fresh medium for another 24 (e) or 48 h (f) incubation. The SC2-P infection was assessed with LUC activity (e), and SARS-CoV-2 infection (supernatant) were evaluated by qPCR (f). Data presented as mean **±** SEM of technical triplicates. ##, **, p<0.01, ###, ***, ∇∇∇, p<0.005 by Student’s *t-*test. ND: not detected. g) The uptake and the degradation of CIPS by macrophages as determined by ICP-MS. Intracellular Indium was used to describe the accumulation and degradation of CIPS and related compounds resulting from CIPS degradation. Differentiated THP-1 cells were treated with 12 pM CIPS up to 24 h (Uptake) and further cultured in fresh medium for 24 and 48 h (Degradation). h) Three-dimensional tomographic images showing CIPS intracellular accumulation after 12 h uptake and 48 h degradation. Images were obtained by soft X-ray transmission microscope (Nano-CT). i-j) Cu chemical transformation and degradation from intracellular CIPS during the uptake and degradation phases, as determined by Cu K-edge XANES. (i) Chemical species of Cu in different reference samples and (j) the speciation of Cu from CIPS during the uptake and the degradation processes. The percentage of Cu forms is reported in Table S2. k-l) CD86 and HLA-DRA gene expression in SARS-CoV-2 treated macrophages in the presence or absence of CIPS. SARS-CoV-2 virus with/without CIPS were incubated with PMA-differentiated THP-1 for 48 h. HLA: HLA-DRA. GAPDH was used as housekeeping gene and the relative gene expression was normalized to the virus infection group. Data presented as mean±SEM (n=3). *, p<0.05, **, p<0.01 by Student’s *t*-test. (m) CIPS induced DC maturation after SC2-P incubation. SC2-P, and SC2-P with CIPS were incubated with DC for 24 h. The CD11c and CD86 positive cells were analyzed by flow cytometer. The double positive (CD11c and CD86) DC were regarded as mature DC cells. Data presented as mean±SEM (n=3). PMA used as positive control. ***, p<0.001 by One ANOVA.

SARS-CoV-2 in macrophages has been reported, although in resting/healthy conditions they do not express ACE2^42^. Since a huge amount of SARS-CoV-2 was internalized by macrophages in the presence of CIPS, we examined whether the intracellular virus was able to infect macrophages. With pseudovirus (SC2-P) or authentic SARS-CoV-2 virus, we show that SARS-CoV-2 uptake through CIPS phagocytosis cannot induce macrophage infection (Fig. 6e–f, SARS-CoV-2 releases in medium) and cause cytopathic or cytotoxic effects (data not shown). This indicates that macrophages still function well with the virus scavenging capacity after massive SARS-COV-2 internalization with CIPS treatment, and avoid viral infection.

We also examined the uptake, accumulation and degradation/clearance of CIPS (12 pM) in macrophages. Based on ICP-MS results for In content, the uptake of CIPS by macrophages was time-depended, and the content of In decreased with time when we removed CIPS from the medium (Fig. 6g). Thus, SARS-CoV-2 that adsorbed on CIPS could be internalized by macrophage together with CIPS, which facilitate SARS-CoV-2 host elimination. Transmission X-ray microscopic imaging (TXM) combined with computer tomography (Nano-CT) was used to visualize intracellular CIPS. Nano-CT results show a substantial accumulation of CIPS within cells after 12 h exposure and likewise strong decreased of intracellular CIPS after 48 h in CIPS-free medium (Fig. 6h). Both the quantitative and imaging results suggest the degradation of CIPS by macrophages. Cu XANES was further used to quantify chemical changes of CIPS during the accumulation in macrophages and its degradation. Cu XANES results show that the speciation of Cu in internalized CIPS is +1 and the chemical bond is Cu-S, respectively. With increasing time, Cu speciation within macrophages changed from +1 to +2, and the chemical form of Cu varied from Cu-S to Cu-O/Cu-OOC- (Fig. 6i–j, Table S1), suggesting that the intracellular form of CIPS changed from oxidation to degradation, most likely within acidic phagolysosomes. These results suggest that CIPS can be progressively degraded and metabolized by macrophages.

The intracellular elimination of SARS-CoV-2 in macrophages that promoted by CIPS, may induce the following immunization process such as antigen presentation. Thus, the expressions of CD86 and HLA-DRA, two molecules important for antigen presentation, were assessed in SARS-CoV-2 treated macrophages and showed greatly up-regulation with CIPS treatment compared to virus alone (Fig. 6k–l). This antigen presenting capacity that promoted by CIPS was further assessed in dendritic cells (DC), and found CIPS was able to increase the population of mature CD11c^+^ CD86^+^ DC in the presence of the SC2-P (Fig. 6m). Thus, the internalization of CIPS by macrophages could facilitate the elimination of the CIPS-trapped SARS-CoV-2, and could further benefit COVID-19 treatment by promoting antigen presentation in the context of MHC-II.

## Discussion

The binding affinity of nAbs with S protein RBD ranges from 0.1 to 100 nM^43,44^, which is similar to the binding affinity between RBD and ACE2 (8.1 nM in our hands, 14.7 nM reported for S protein binding to ACE2)^40^. The comparable affinity suggests that only very few of the currently available nAbs may have sufficient affinity for RBD to efficiently compete with its binding to ACE2 thereby inhibiting cell infection. Here, we have identified the 2D CIPS NS as an effective nano-viscous material able to capture the S protein and inhibit cell infection by SARS-CoV-2 (Fig. 3–5). CIPS strongly binds to the S protein of SARS-CoV-2 and interferes with the ability of the S protein RBD to bind to host ACE2 (Fig. 4). The binding affinity of CIPS for the S protein RBD is <1 pM, suggesting the formation of very stable complexes and highly efficient inhibition of the virus interaction with ACE2. Computational simulations are consistent with experimental data (Fig. 5, fig. S3 and S12), and show that CIPS/RBD exhibits the lowest interaction energy and the strongest electrostatic attraction (Fig. 5a, d, fig S18, Table S4). The effective binding of CIPS with 6 out of the 11 ACE2-binding amino acid residues of the SARS-CoV-2 RBD accounts for the efficient inhibition of viral infectivity displayed by CIPS (Table 1). The strong RBD binding affinity displayed by CIPS and its capacity to inhibit SARS-CoV-2 infection are apparently selective and are maintained in complex bio-systems. Although the Asn501 is replaced by Tyr501 in the new SARS-CoV-2 variant, the N501Y RBD variant could be still effective captured by CIPS, suggesting a consistent anti-SARS-CoV-2 capacity of CIPS (fig. S16). The similarity between the SARS and SARS-CoV-2 RBDs and infection mechanisms suggests that CIPS is potentially useful also for containing the SARS virus infection and spreading (fig. S19).

CIPS proved safe and biocompatible both *in vitro* (fig. S5) and *in vivo* (fig. S20), showing quick degradation and removal by macrophages (Fig. 6 g–j). Importantly, the SARS-CoV-2 virus adsorbed on CIPS can be efficiently taken up by macrophages, shuttled to phagolysosomes and completely degraded (Fig. 6 a–d). This suggests that CIPS could be successfully used to capture the virus in the body and direct it to degradation by facilitating its elimination by macrophages. Another important consequence of the phagocytic virus uptake by macrophages and its intracellular degradation through the endocytic/lysosome pathway, caused by CIPS, is the expected generation of viral peptides to be presented in the context of MHC-II molecules, *i.e.*, the pathway of antigen presentation that induces a strong antibody response (Fig. 6 k-m). Thus, the capacity of CIPS to direct the virus to macrophages has the double advantage of promoting viral destruction (thereby limiting infection) while at the same time triggering anti-viral immunization.

Thus, CIPS is a safe, biocompatible, biodegradable 2D NM capable of inhibiting the infection and promoting the elimination of SARS-CoV-2. The binding between CIPS and the SARS-CoV-2 S protein RBD (K_D_ <1 pM) is 10,000x stronger than the affinity of the virus for ACE2, suggesting that the CIPS-captured virus will not be released and will not infect cells. Notably, CIPS binding is 100x stronger than that described for the best nAbs. In addition, CIPS promotes SARS-CoV-2 uptake and degradation by macrophages through a pathway that promotes virus-specific antibody production. Eventually, its glue-like selective binding capacity makes CIPS applicable both as disinfection agent and in surface coatings, and as a nanodrug candidate for treating COVID-19 (Fig. 7).

**Fig. 7.**
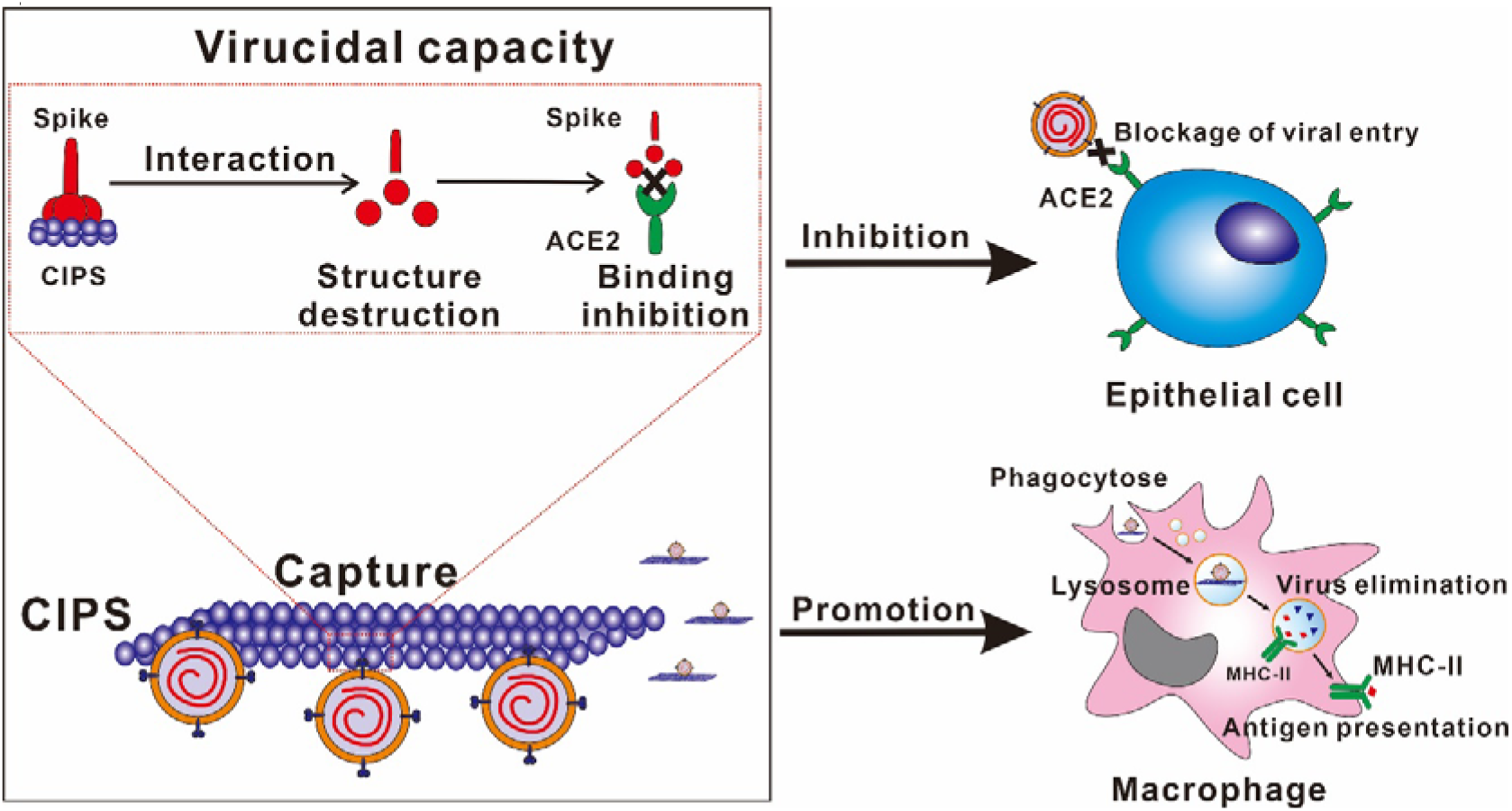
The anti-SARS-CoV-2 capacity of CIPS “nano-glue”. The mechanism that possessed by CIPS on SARS-CoV-2 restraining and elimination. The RBD of S protein can be tightly absorbed to CIPS that induced RBD deformation. The selectively binding capacity of CIPS on RBD with a *K*_D_ < 1 pM that is 1000 times higher than the reported neutralizing antibodies, causes the inhibition of SARS-CoV-2 infection. Further, the CIPS-trapped SARS-CoV-2 can be effectively phagocytized by macrophages and lead to virus elimination in lysosome, which further facilitates COVID-19 treatment.

## Materials and Methods

### Preparation of CIPS nanosheets

Bulk CuInP_2_S_6_ (CIPS) material was synthesized by solid state reaction according to previous publication.^45^ CIPS nanosheets (NSs) were exfoliated from the bulk CIPS crystals with Li-intercalation by using n-butyl lithium assisted by ultrasound sonication. After the exfoliation, NSs were centrifuged at 1500 g for 20 min and then rinsed with H_2_O twice to remove free n-butyl lithium in the suspension. The collected suspension was finally lyophilized and stored in dry condition at −20ºC.

### CIPS effects on SARS-CoV-2 virus with Vero-E6 cells

The experiments were performed in a biosafety level 3 laboratory with standard regulations. SARS-CoV-2 was isolated in the laboratory and propagated on Vero-E6 cells. Upon infection, Vero-E6 cells were seeded in a 96-well plate at a density of 2×10^4^ cells/well, and cultured overnight. CIPS at various concentrations (1.5-96 pM) and the SARS-CoV-2 (100TCID50) were added together to Vero-E6 cells and incubated for 2 h. Cells were then washed twice with warm 1x PBS and cultured for additional 48 h with fresh medium or CIPS-containing medium. Non-infected cells and CIPS alone were used as controls. Supernatants were collected after 48 h incubation, and the SARS-CoV-2 was quantitatively analysed by real-time PCR with a commercial COVID19 detection kit (DAAN Gene Ltd., Guangzhou, China. Cat# DA0931) targeting the nucleocapsid protein (N) and the open reading frame 1ab (ORF1ab/RaRp). In brief, the N and ORF1ab were detected using FAM and VIC labelled Taqman probe by Applied Biosystems^™^ 7500 (ThermoFisher Scientific). The amplification was performed as follows: 50°C for 15 min, 95°C for 15 min, followed for 45 cycles with 94°C for 15 sec and 55°C for 45 sec. For quantitative evaluation, plasmid standards were used to establish a standard curve with serial dilutions (copy number *vs.* fluorometric value), and used to calculate the RNA copy number of SARS-CoV-2 from the fluorometric value of the samples. The standard curve is y=50×10^(12.352-0.2771*C_T_). The infection rate was calculated and normalized to virus infection controls. The conditions of Vero-E6 cells after SARS-CoV-2 infection were observed at 48 h, and the optical microscopy images were acquired. The occupied space of the cells was analyzed by ImageJ and shown as percentage of the area fraction covered by cells, as an estimate of intact healthy cells in the image.

### Antiviral effect of CIPS on Vero-E6 cells with SARS-CoV-2 post-infection

Vero-E6 cells were seeded in 24-well plate overnight with a density of 5×10^4^ cells/well. Cells were challenged with SARS-CoV-2 (12000 pfu) for 2 h, and the uninfected virus were washed out by PBS. Cells were continuously cultured in fresh medium or medium containing CIPS with various concentrations (24, 48, 96 pM) for another 48 h. Cells were collected and lysed with RNAplus (TAKARA, Japan). Intracellular SARS-CoV-2 was evaluated by qPCR by targeting S protein. Cellular 18S was used as housekeeping gene to normalize the intracellular number of viruses as virus per cell.

### Human airway tissue cultures

Human nasal differentiated epithelial Air-Liquid-Interface (ALI) cultures were derived from patient bioptic tissue. The nasal epithelium was obtained from surgically excised rhinopolyp tissue. The tissue was washed with sterile PBS, the healthy epithelium portion of the tissue dissected and cut into small pieces with sterile scissors. The tissue pieces were dissociated with Dispase I (STEMCELL Technologies, Vancouver, Canada) and centrifuged to collect dissociated cells, Cells were plated in culture dishes with a feeder of 3T3 fibroblasts, and expanded using PneumaCultTM-Ex Plus Medium (STEMCELL Technologies). After progenitor cell expansion, the cultured cells were transferred onto 0.4 μm membranes of 24-transwelll-plates, and differentiated in ALI culture conditions with PneumaCultTM-ALI Medium (STEMCELL Technologies). All the human tissue-related work was approved by the ethics committee of the Seventh Affiliated Hospital of Sun Yat-sen University, Shenzhen (No. 0720) and the ethics committee of Shenzhen Institutes of Advanced Technology, Chinese Academy of Sciences (SIAT-IRB-200215-H0415). Written informed consents were obtained from all participating patients.

### CIPS effects on SARS-CoV-2 virus infection of human airway epithelial tissue cultures

A SARS-CoV-2 inoculum of 10000 pfu per 50 μl was added to a single airway epithelial tissue-bearing transwell insert placed in a round-bottom 24-well plate (well diameter 0.33 cm^2^) with/without 24 pM CIPS and incubated at 37°C for 1 hour. The inoculum was removed, and tissue-bearing inserts washed 3x with PBS to remove unbound virus. Each virus-infected tissue-bearing insert was transferred to a single well of a new 24-well plate (Corning, USA) with 500 μl of fresh serum-free growth medium (05001, STEMCELL Technologies) or CIPS-containing medium, and incubated at 37°C with 5% CO_2_ for 24 h. To analyze the virus infection, 150 μl Trizol (Qiagen, Hilden, Germany) were added onto the upper surface of epithelial culture. Viral RNA in the collected samples was determined by RT-qPCR with a SARS-CoV-2 diagnostic kit approved by the China CDC (BioGerm, 2C-HX-201-2), following the manufacturer’s protocol. To examine the histological changes caused by SARS-CoV-2 infection and CIPS treatment, paraffin-embedded epithelia were sectioned at a thickness of 3 μM and examined after hematoxylin and eosin (H&E) staining.

### CIPS captures SARS-CoV-2 virus

In a P3 laboratory, the SARS-CoV-2 (6000 pfu) were incubated with CIPS (6 and 12 pM in 200 μl PBS) for 2h, and centrifuged (3000 rpm for 5 min) to separate the CIPS-trapped SARS-CoV-2 in precipitates. The RNA of the SARS-CoV-2 was extracted for reverse transcription (TAKARA, Japan). The SARS-CoV-2 was quantitative analyzed with real time PCR by targeting S protein.

### Evaluation of binding affinity between CIPS and proteins

Biolayer Interferometry (BLI) technique was used to measure the binding affinity of CIPS or CIPS-RBD complex to proteins, using an Octet RED96e system (FortéBio, Bohemia, NY, USA). For the interaction between CIPS and proteins, the running buffer was a mixture of 0.1% BSA and PBS-P (10 mM phosphate salts, 2.7 mM KCl, 137 mM NaCl and 0.05% P20 surfactant). CIPS NSs at 12 pM were immobilized on the surface of AR2G sensor chip (immobilization time 600 s and immobilization height 3.5 nm). The sensor was then immersed into the solution of different proteins, the receptor binding domain of the SARS-CoV-2 spike protein (RBD, from DIMA Biotechnology Ltd, China), human serum albumin (HSA, from Solarbio Science & Technology Co., Ltd. China), fibrinogen (FG, from Sigma-Aldrich), immunoglobulin G (IgG, from Sigma-Aldrich), transferrin (Tf, from Sigma-Aldrich), high density lipoprotein (HDL, from Yiyuan biotechnology Ltd, Guangzhou, China)) at different concentrations (12.5-200 nM) and then their binding was assessed with an association time of 300 s and a dissociation time of 600 s. The same procedure was adopted to evaluate the binding affinity of graphene oxide (GO) and MoS_2_ NSs to RBD. For their immobilization, the concentrations for GO and MoS_2_ were 20 μg/ml and 2 mg/ml, respectively.

To study the binding affinity of RBD to ACE2 (DIMA Biotechnology LTD), RBD was immobilized on the surface of the AR2G sensor chip using the amine-coupling procedure (immobilization time 300 s), and binding to ACE2 assessed with an association time of 300 s and a dissociation time of 600 s.

To assess the interference of CIPS with the RBD binding to ACE2, 200 μl of CIPS (12 pM) dispersed in the above running buffer were added to a 96-well microplate. CIPS NSs were immobilized on the surface of the AR2G sensor chip with the same parameters mentioned above and then immersed into the solution of RBD at 50 nM, and binding assessed with an association time of 300 s and a dissociation time of 600 s. Next, blocking with BSA was performed by adding 0.1% BSA into running buffer for ACE2 for 1200 s. Furthermore, the sensor was immersed into ACE2 solutions at different concentrations (12.5-200 nM) with an association time of 300 s and a dissociation time of 600 s. The data were collected and analyzed with the Data Analysis 11.0 Software. The affinity constants (K_D_), association rate constants (K_on_) and dissociation rate constants (K_off_) were calculated by fitting the curves using the 1:1 kinetic binding model.

### Determination of secondary structure of proteins by Circular Dichroism

The secondary structure of RBD in the presence or absence of CIPS was assessed by Circular Dichroism (CD) (JASCO, J-810). RBD (500 μl at 200 μg/ml in 0.01 M phosphate buffer pH 7.4) was mixed with CIPS (15, 30, and 60 pM) and then added into the liquid well with 1 mm thickness. CD spectra were collected between 190 and 250 nm. Each sample was measured six times and the averaged spectra were obtained. Spectral data were processed with the CD tool software (available at http://cdtools.cryst.bbk.ac.uk). The baseline (between 235 and 240 nm) was subtracted. Normalized data were analyzed by the web server DICROWEB to calculate the ratio of secondary structure, and the smoothed data are shown.

### CIPS effects on SC2-P elimisnation and infection of THP-1 differentiated macrophages

The THP-1 cells were seeded in a 24-well plate with cover slide at a density of 5×10^4^ cell/well (for elimination), or in a 96-well plate at a density of 1×10^4^ cell/well (for infection), and treated with 100 ng/ml PMA 24 h for macrophage differentiation.

For elimination experiment, CIPS (12 pM) was used to pre-incubate with SC2-P for 2 h. The SC2-P with/without CIPS pre-treatment was added to the PMA differentiated THP-1 cells for 24 h, named as “phagocytosis”. Here after, fresh medium was used to replace the SC2-P containing medium and cultured for another 24 h, named as “degradation”. Bafilomycin (BM), the lysosome inhibitor, was added to macrophages 6 h before the end point of each experiment. The macrophages phagocytosis and degradation of SC2-P were detected by IF.

For infection experiment, THP-1 cells were co-cultured with SC2-P (2×10^5^ copies) for 24 h in the presence or absence of CIPS (24 pM), and then washed by warm DMEM to remove extracellular virus. Cells were cultured with DMEM for additional 24 h. BM (100 nM) was added at 18 h. Cells were lysed for luciferase evaluation (E1910, Promega, USA).

### CIPS effects on SARS-CoV-2 virus elimination and infection of THP-1 differentiated macrophages

The experiments were performed in a biosafety level 3 laboratory with standard regulations. The THP-1 cells were seeded in a 24-well plate at a density of 5×10^4^ cell/well, and treated with 100 ng/ml PMA 24 h for macrophage differentiation.

For elimination experiment, SARS-CoV-2 (12,000 pfu) were added to macrophages for 4 h (phagocytosis). Cells were then washed twice with warm 1x PBS and cultured for additional 48 h with fresh medium (degradation). Macrophages were collected for RNA extraction. The intracellular SARS-CoV-2 was quantitative analyzed by real time PCR by targeting S protein.

As for CIPS effect, authentic SARS-CoV-2 virus (12,000 pfu) was pre-incubated with CIPS (12 pM) for 2h. The SARS-CoV-2 with/without CIPS pre-treatment was added to the PMA differentiated THP-1 cells for 48 h in the presence or absence of BM (100 nM). Macrophages were washed twice with PBS and subjected to RNA extraction by RNAplus (TAKARA, Japan). The intracellular SARS-CoV-2 was quantitative analyzed by real time PCR by targeting S protein. Gene expression of CD86 and HLA-DRA was also quantitative analyzed.

For infection experiment, CIPS (12 pM) was used to pre-incubate with authentic SARS-CoV-2 virus (12,000 pfu) for 2h. Macrophages were treated by SARS-CoV-2 (with/without CIPS) for 4 h, and were then washed twice with warm 1x PBS. Macrophages were cultured in fresh medium with/without BM (100 nM) for additional 48 h. Supernatants were collected, and the SARS-CoV-2 was quantitatively analyzed by real-time PCR with a commercial COVID19 detection kit (DAAN Gene Ltd., Guangzhou, China. Cat# DA0931) targeting the nucleocapsid protein (N) and the open reading frame 1ab (ORF1ab/RaRp)

### Statistical analysis

Results are presented as mean values of replicate experiments or replicate samples within one best representative experiment, as indicated in the Figure legends. Statistically significant differences (*p*<0.05) were determined by the Student’s *t*-test or ANOVA analysis where appropriate.

## Acknowledgements

This work was financially supported by the National Natural Science Foundation of China (31701005 and 31971322), Shenzhen Science and Technology Program (GJHZ20190821155803877), CAS President’s International Fellowship Initiative (PIFI, 2020VBA0028), the National Basic Research Program of China (2016YFA0201600, 2016YFA0203200, 2020YFA0710702), and the Users with Excellence Project of the Hefei Science Center CAS (2018HSC-UE004), and was supported by the State Key Laboratory of Natural and Biomimetic Drugs, Peking University.

## Author Contributions

Y.L. and L.W. conceived and designed the project. Y.L., L.W., C.C., and H.L. supervised the study. G.Z. and G.C. performed the biology and biochemistry experiments. H.L. and K.L. designed and validated the pseudovirus systems. L.L., J.Q., S.L., and Y.F. performed SARS-CoV-2 viral experiments in the P3 lab. Y.C., L.W., J.W., and Y.F.L. did BLI, CD, Nano-CT, and XAFS experiments. W.X. analyzed EXAFS data. G.Z., G.C., and Y.C did ICP-MS experiment. L.W. took part in MD simulation design and discussion with Boyubio Company. P.Y. prepared and characterized CIPS NSs. Q.S. did the *in vivo* safety experiment, Q.S. and G.W. contributed to the synthesis and characterization of nanomaterials. Y.L., L.W., C.C. H.L., D. B., Y.Z. G.Z., L.F., and Y.C. analyzed and discussed the data. Y.L. D. B., G.Z., and L.W. wrote the manuscript. #These authors contributed equally.

