## Supplementary material for "Spike Protein Targeting “Nano-Glue” that Captures and Promotes SARS-CoV-2 Elimination": fig.S

### Materials and Methods

#### Characteristics of CIPS NSs

A 20 μl aliquot of CIPS NSs suspension was dropped onto a lacey carbon TEM grid and then observed under a TEM (Tecnai G2 F20 S-Twin, FEI, USA). The powder X-ray diffraction (XRD) patterns were recorded on a Bruker AXS D8 ADVANCE diffractometer (Bruker AXS GmbH, Karlsruhe, German) with Cu Kα radiation. A 50 μl aliquot of CIPS NSs suspension was dropped on the smooth mica substrate and dried by the nitrogen gas. Samples were then observed under an AFM (Veeco Instruments Inc., USA) to measure size distribution and thickness of the CIPS flakes.

#### Chemical speciation and coordination structure analysis for elements in CIPS NS

X-ray absorption fine structure (XAFS), including X-ray absorption near-edge structure (XANES) and extended X-ray absorption fine structure (EXAFS), was used to characterize chemical species of Cu, In, P, and S, as well as the coordination structure of Cu, in CIPS nanosheets. Cu K-edge XANES was used to assess the changes in the Cu chemical form during the uptake of CIPS by THP-1 derived macrophages, intracellular accumulation and degradation of CIPS. Cells, in complete medium containing 10% FBS, were treated with 12 pM CIPS for 12 h and then cultured in fresh medium without CIPS for 24 h. After detachment from culture dishes with trypsin, cells were rinsed with PBS, pelleted by centrifugation, lyophilized, and finally transferred to a tube filled with nitrogen gas before XAFS measurement.

XAFS spectra for Cu K-edge and In L_3_-edge were collected at the beamline of BL-14W1 in Shanghai Synchrotron Radiation Facility (SSRF) and the beamline of 1W1B in Beijing Synchrotron Radiation Facility (BSRF), China. Transmission mode was adopted to collect the XANES for both CIPS and the reference samples of Cu and In. Fluorescence mode was used to determine Cu K-edge XANES for CIPS within cells. In addition, S K-edge XANES spectra were collected at the beamline of 4B7A in BSRF and were used to quantify chemical form of S in reference samples and CIPS under the total electron yield mode. Afterwards, XAFS data were normalized and analyzed with least-squares fitting (LSF) to calculate the ratio of various species for interested elements using the IFEFFIT Athena software (CARS, the Consortium for Advanced Radiation Sources at University of Chicago, IL, USA). The same software was used to analyze the coordination information of Cu in the CIPS samples.

#### Cell lines and cell culture

The human kidney cell line HEK293T (293T), ACE2-GFP stably transfected HEK293T (ACE2/293T), the human bronchial epithelial cell line 16HBE (Cell Bank of Peking Union Medical College, China), the human type II-like lung carcinoma cell line A549, primary human adult epidermal keratinocytes (pHEK-Ad; Lonza, Basel, Switzerland), murine RAW264.7 cells and African green monkey Vero-E6 cells were all cultured in DMEM (HyClone, Logan, UT, USA) with 10% fetal bovine serum (FBS; Gibco, Thermo Fisher Scientific, Inc., Waltham, MA, USA) and 1% penicillin/streptomycin solution (P/S, stock concentration as 1×10^4^ units/ml of penicillin and 10 mg/ml streptomycin; HyClone) at 37°C in humidified atmosphere with 5% CO_2_. The human monocytic leukemia cell line THP-1 was cultured in 1640 RPMI medium (HyClone) containing 10% FBS and 1% penicillin/streptomycin solution.

#### Development of HEK-293T cell line with stable expression of ACE2-GFP

The coding sequence of ACE2-GFP was inserted into the multiple cloning site of pCDH vector (pCDH-EF1-MCS-IRES-Puro, CD532A-2, System Biosciences) with *Eco*RI (Thermo Fisher Scientific, Inc.). By using lentiviral transfection system, the ACE2-GFP vector was transferred into HEK293T. Stably transfected 293T cells (ACE2/293T) were selected with puromycin (Solarbio, Beijing, China) and confirmed by confocal microscopy.

#### Cell viability assay

Cells (1x10^4^ cells/well of 96-well plates) were cultured overnight and then exposed to different NMs including CIPS (1.5-96 pM; 6 replicates) for 24-48 h. Supernatant was then replaced with fresh DMEM with 10% CCK8 solution (transformed in formazan by enzymes of living cells) and incubated for one additional hour at 37°C. Formazan absorbance at 450 nm was measured with a Microplate Reader (Multiskan GO, Thermo Fisher Scientific, Inc.). The cell viability and cytotoxicity were calculated, normalized to the controls and presented as percentage.

#### Plasmid construction and the establishment of pseudovirus

***Plasmid construction***

The full-length coding sequence of the SARS-CoV-2 Spike glycoprotein, SARS-CoV Spike glycoprotein and VSV-G were fused with a Flag tag and inserted into the pCAGGS vector by *Eco*RI (Thermo Fisher Scientific, Inc.) to obtain the plasmids of pCAGGS-SC2-S-Flag, pCAGGS-SARS-S-Flag and pCAGGS-VSV-G-Flag, respectively.

***Pseudovirus establishment***

The SARS-CoV-2 pseudovirus (SC2-P), SARS pseudovirus (SARS-P) and VSV-G pseudovirus (VSV-G) were produced with the method previously reported^1^. In brief, the pCDH-LUC (luciferase gene), the packaging plasmid psPAX2 (#12260, Addgene) and the constructed plasmids were transferred to HEK293T cells by polyetherimide (PEI). After 48 or 64 h, the pseudovirus-containing cell culture medium was collected, centrifuged at 1000 g for 5 min and purified with a 0.45 μm filter to remove cell debris. The pseudovirus-containing medium was further added to the top of 25% sucrose and centrifuged for 2 h at 100,000 g at 4°C to concentrate and obtain the purified pseudovirus.

***Pseudovirus identification***

The constructed plasmids were transfected in HEK293T cells with Lipofectamine 2000 (Thermo Fisher Scientific, Inc.) for 24 h. Cells then were lysed with a protease inhibitor cocktail (Roche, Basel, Switzerland) containing RIPA buffer (Beyotime Biotechnology, Shanghai, China), and centrifuged 12000 rpm for 5 min at 4°C. The supernatants were boiled with loading buffer at 100°C for 5 min, run with 8% SDS-PAGE and immunoblotted with anti-Flag (AF0036, Beyotime Biotechnology, Shanghai, China). To detect S protein incorporated into pseudovirus, the purified pseudoviruses were boiled with loading buffer at 100°C for 5 min and run with 8% SDS-PAGE and immunoblotted with anti-Flag. The blots were analyzed with an Amersham Imager AI600 (GE, Boston, MA, USA).

#### ACE2 knockdown in Vero-E6 cells and validation of pseudovirus infectivity

A commercially synthesized siRNA was transfected with Lipofectamine 2000 (Thermo Fisher Scientific, Inc.) into Vero-E6 cells for endogenous ACE2 knockdown. ACE2-siRNA forward sequence: CACGAAGCCGAAGACCTGTTCdTdT, ACE2-siRNA reverse sequence: dTdTGAACAGGTCTTCGGCTTCGTG. After 48 h, cells were collected for immunoblot analysis with anti-ACE2 (sc-390851, Santa Cruz Biotechology, Dallas, TX, USA).

The various pseudoviruses were incubated with Vero-E6 cells (with/without ACE2 knockdown), and infection identified with immunofluorescent (IF) anti-Flag antibody using a confocal laser scanning microscope (Leica SD AF, Wetzlar, Germany). Mean fluorescent intensity (MFI) was quantitative analysis of the acquired images by ImageJ.

#### Evaluation of nanomaterials and CIPS inhibition of SARS-CoV-2 pseudovirus infection with luciferase assay

ACE2/293T or Vero-E6 cells were seeded (1x10^4^ cells/well) in a 96-well plate and cultured overnight. LUC-carrying SC2-P (2x10^5^ copies) was pre-incubated with different NMs for 2 h at RT and the mixtures were then added to cells for 2 h. Cells were then washed by warm DMEM to remove extracellular virus, and cultured with DMEM for additional 40 h. Cells were lysed with 50 μl 1x lysis buffer (E1910, Promega, USA) and shaking for 5 min at RT. The cell lysate (30 μl) was mixed with 20 μl luciferase substrate, and the luciferase activity was immediately measured with a microporous plate luminescence detector (Glomax 96, Promega, USA) and expressed as relative luciferase units (RLU). Experiments were run in triplicate, and the RLU were analyzed for each biological sample with three technical replicates. The SC2-P infection rate was calculated and normalized to controls. For NM screening, only ACE2/293T cells were used, while Vero-E6 cells were also examined for further analysis of CIPS activity. The possible CIPS interference on luciferase activity was assessed by adding CIPS (0.048 pmol) to the medium with a known amount of luciferase and incubated for 2 min. The luciferase activity was then detected and compared with no CIPS control, and showed no interference of CIPS on luciferase activity.

#### Immunofluorescence evaluation of SARS-CoV-2 pseudovirus infection

HEK293T cells were seeded at 4x10^4^ cells/well of a 24-well plate, each containing a 14 mm coverslip, cultured overnight and transfected with ACE2-GFP for 24 h to generate ACE2 overexpressing (ACE2-OE) cells.

SC2-P (4×10^6^ copies) was pre-incubated with different CIPS concentrations for 2 h at RT. The pre-incubated SC2-P and CIPS were then added to ACE2/293T or Vero-E6 cells, which were further cultured for different times (0.25, 0.5, 1, 2 and 15 h) at 37^o^C. Cells were then fixed with 4% paraformaldehyde and incubated with antibodies anti-Flag (AF0036, Beyotime Biotechnology) and anti-GFP (sc-9996, Santa Cruz Biotechnology) for 2 h, and further with goat anti-rabbit IgG Alexa Fluor 555 (A21429, Thermo Fisher Scientific, Inc.) and goat anti-Mouse IgG Alexa Fluor 488 (A11029, Thermo Fisher Scientific Inc.) antibodies for one addition hour. The SC2-P infection was examined as fluorescent signal with a confocal laser scanning microscope (Leica SD AF). Red dots in the IF images (representing individual SC2-P) were counted for each cell (cells ≥ 3), and the mean fluorescent intensity (MFI) of the IF image was analyzed by ImageJ.

#### Infectivity of SARS-CoV-2 pseudovirus exposed to CIPS in complex mixtures

SC2-P was admixed with FBS or BSA in volume ratios SC2-P to FBS/BSA of 1:1 to 1:10, and then incubated with 12 pmol for 2 h. The mixture was washed with 1 ml PBS and centrifuged at 4000 rpm for 10 min at RT. The S protein absorbed on CIPS was detected in WB with anti-Flag antibody. To assess the pseudovirus infectivity in these conditions, ACE2/293T cells were exposed to the mixture of SC2-P, FBS/BSA and CIPS described above and incubated for 2 h. The infected ACE2/293T cells were cultured for 40 h and lysed. Luciferase activity, representing the SC2-P infection, was assessed with a luminescence detector (Glomax 96, Promega, USA).

#### The binding analysis of CIPS with SC2-P and spike protein

The SC2-P (4×10^6^ copies) was pre-incubated with CIPS for 2 h at RT. The mixture was added with 1 ml PBS and centrifuged 4000 rpm for 5 min to get the precipitate, by which to detect the TEM, UV-VIS, Z-potential and Western Blot (WB).

For the S protein binding analysis, the constructed plasmid of pCAGGS-SC2-S-Flag was transfected in HEK293T cells with Lipofectamine 2000 (Thermo Fisher Scientific, Inc.) for 24 h. Cells then were lysed with a protease inhibitor cocktail (Roche, Basel, Switzerland) containing RIPA buffer (Beyotime Biotechnology, Shanghai, China), and centrifuged 12000 rpm for 5 min at 4°C to obtain the S protein in supernatant. The obtained S protein containing supernatant was incubated with CIPS for 2 h at 4°C and subjected to centrifugation for 4000 rpm, 4°C for 5 min. The obtained precipitate was used to detect the binding of CIPS with S protein by WB with anti-Flag (AF0036, Beyotime Biotechnology, Shanghai, China).

#### Molecular dynamics (MD) simulation system

The structure and topology of CIPS is reported in previous publications^2,3^. The atomic structure of CIPS exhibits the monoclinic symmetry (system) with space group Cc. The lattice parameters are a = 6.095 Å, b = 10.564 Å, c= 13.623 Å and α = 90.000°, β = 107.101°, γ = 90.000°. Three S atoms in a triangular pattern connected by a P atom form a plane, other three S atoms and a P atom form a bottom plane opposite to the former one. These two planes are connected by the two P atoms and the In atom inlaid in the middle. In the plane projection direction, Cu and In atoms are at the vertex of the long diagonal of the parallelogram. The distribution of Cu atoms in the two planes is heterogeneous, with 15% Cu atoms in the upper plane and 85% in the lower plane. These two planes, forming a layer of CIPS, are 3.354 Å apart and are connected by van der Waals (vdW) forces. The vertical distance between two layers is about 3.2 Å.

In the molecular dynamics simulation, the partial charge of each atom is a key parameter. The electrostatic charges for the CIPS were computed based on the MK-RESP (Merz-Kollman Restrained Electrostatic Potential) methodology^4,5^ using the Multiwfn code^6^, and the equivalence and valence constraint were taken into account. The vdW parameters were obtained from Universal Force Field^7^.

The atomic structure of the RBD of the SARS-CoV-2 S glycoprotein was extracted from the crystal structure of S protein binds ACE2 complex (PDB ID 6M17, chain F). This region contains 183 residues (from 336 to 518). The conformation of RBD could be approximated to a right triangular prism. There are 5 β strands (β1 to β4 and β7) in the middle with 3 α helices (α1 to α3) in one side and another 2 helices (α4, α5) in the other side, the rest 2 β strands (β5, β6) formed a right-angle side. The protein surface was then divided into 5 sides. The initial configurations of the simulation were constructed with each side facing the CIPS surface. Because of CIPS heterogeneity, both surfaces were considered. In the simulation box, CIPS was placed along the x-y plane, and the size was 73.152 ×52.826 Å^2^. RBD (in five different orientations corresponding to the 5 identified sides of the molecule) was placed 5 Å above the CIPS surface to ensure that there was no contact in the initial state. The height of the simulation box was from 80 to 100 Å varying with different sides of RBD. The charmm27 force field^8^ was used to describe the interaction of RBD with CIPS using UFF parameters. The TIP3P water model^9^ and Na^+^, Cl^−^ ions were added to neutralize the system and maintain a physiological concentration of 0.15 M.

The GROMACS software package (version 5.1.5) was used to perform molecular dynamics simulations^10^. The complete system was firstly minimized to remove unfavorable geometric clashes and then equilibrated for 2 ns in the NVT ensemble. During the equilibration simulation, the positions of protein Cα atoms were restrained (1000 kJ mol^−1^ nm^−2^) while maintaining the CIPS atoms frozen, which allowed us to use a time step of 2 fs. The temperature during this procedure was controlled at 300 K in the Nose-Hoover thermostat with a collision frequency of 0.5 ps^−1^.^11^ Periodic boundary conditions (PBC) were set for all directions. The vdW interactions were computed with a cutoff distance of 12 Å, while long-range electrostatic interactions were handled with the particle mesh Ewald (PME) method^12^. Water molecules were constrained by the SETTLE algorithm^13^, and solute hydrogen bonds were constrained to their equilibrium values employing the LINCS algorithm^14^. The coordinates were saved every 100 ps and all simulation snapshots were rendered with the program VMD^15^. Afterwards, we initiated the simulation for 100 ns in the NPT ensemble using the Parrinello-Rahman method^16^. The pressure was maintained at 1 atm by coupling the semi-isotropic (X+Y, Z) directions of the system with the time constant of 5 ps^-1^. The protein atoms were free to move while maintaining the positional restraints (1000 kJ mol^−1^ nm^−2^) on the CIPS atoms. The other settings were the same as the equilibration simulations. To determine the adsorption surface of CIPS, 6 trial runs with the same RBD side were performed, three of them facing the upper CIPS surface (containing 15% Cu) and the other three facing the lower surface (containing 85% Cu). Then, five independent runs for each side of RBD facing the selected CIPS surface. Lastly, the final configurations obtained in each run (25 configurations totally) were put together for clustering.

For the GO system, the GO NS was constructed by modifying the surface of graphene with oxide, hydroxyl (-OH), and epoxy (=O) groups. These oxidation groups were uniform randomly distributed on both the upper and lower surfaces of graphene. The ratio of -OH to =O was 1:1 and the GO sheets have a C/O ratio of 3/1. The partial charge and Lenard-Jones parameters for atoms in epoxy and hydroxyl groups of GO are listed in extended data Table S6. The non-oxidized carbon atoms of the GO nanosheet were modeled as uncharged Lenard-Jones particles with a cross section of σ_cc_ = 0.34 nm and a potential well depth of ε_cc_ = 0.3598 kJ/mol^17^. The size of GO sheet was 81.219 × 62.984 Å^2^ that provides enough area for RBD adsorption.

For the molybdenum sulfide (MoS_2_) NS, we used a trigonal prismatic (2H) stacking pattern corresponding to the P63/mmc space group^18^. Only one slab of MoS_2_ with the top-view hexagonal atomic arrangements contains molybdenum atom sandwiched between two sub-layers of the sulfur placed alone x-y plane. The size of MoS_2_ sheet 1s 86.116 × 74.654 Å^2^, which provides sufficient space for RBD adsorption.

#### Quantitative analysis for cellular uptake of CIPS

PMA differentiated THP-1 cells, which developed macrophage-like morphology and functions, were seeded in a 6 well-plate at a density of 1x10^6^ cells/well and cultured overnight. Cells were treated with CIPS (12 pM) for 2, 3, 6, 12 and 24 h for uptake studies, and then cultured in fresh medium for additional 24 and 48 h for degradation study. After rinsing 3 times with PBS, cells were collected, counted, and centrifuged. HNO_3_ was added to cell pellets overnight before transfer to conical flasks that were placed onto a hot plate 3 h at 150°C, to allow ion release. Samples were cooled to RT and diluted with 2% HNO_3_ solution to a volume of 3 ml. A Yttrium solution (10 ng/ml in 2% HNO_3_) was used as internal standard. A series of Cu dilutions (0.1-500 ng/ml in 2% HNO_3_) were prepared as standard. All solutions were measured at least three times by ICP-MS to obtain the averaged results.

#### Nano-CT imaging for the accumulation and degradation of CIPS

Single cell imaging was performed to observe the accumulation and the degradation of CIPS within the THP-1 differentiated macrophages. We used cryo-soft X-ray transmission microscope (TXM) (nano-CT) at the beamline BL07W of the National Synchrotron Radiation Laboratory (NSRL, Hefei, China) to observe the samples. Cells were firstly differentiated on the non-carbon formvar of a nickle 100 mesh with medium containing 100 ng/ml PMA overnight and then treated with 12 pM CIPS for 12 h (uptake samples). Then, cells on the mesh were placed in fresh medium for another 24 and 48 h (degradation samples). After PBS rinsing and cell fixation with 10% paraformaldehyde for 20 min, the nickle grid was immersed in liquid ethane and then inserted into a sample holder in liquid nitrogen. The sample holder was transferred to the chamber of a TXM. Under an energy of 520 eV, the soft X-ray beam focused onto the cells and a zone plate was used as an objective to magnify the cell onto a CCD camera with a 13 μm field of view and a 30 nm spatial resolution. To observe cells by a TXM, cells were rotated from -60° to +60° and a series of projected images were collected for a continuous 121 times with 1-degree intervals and 2-s exposure time. We then aligned the tilt series using XMController, performed 3D tomographic reconstruction by XMReconstruction, and visualized and segmented cell structures such as organelles, cytoplasmic membrane, nucleus *via* Amira (FEI, USA). Organelles, cytoplasmic membrane, CIPS-containing organelles, and the aggregated CIPS were defined based on the difference in their linear absorption coefficient. Because the cell size is larger than the view field, the upper or the bottom part of the cells was chosen for 3D imaging. To visualize cellular components and CIPS, we labeled the cytoplasm in cyan, the CIPS-containing organelles in blue, CIPS in red, and the nucleus in pink.

#### Safety assessment of CIPS *in vivo*

Animal experiments were performed under the Guide for Care and Use of Laboratory Animals and were approved by the Institutional Animal Care and Use Committee (IACUC) of the Shenzhen Institutes of Advanced Technology (SIAT). C57BL/6J mice were intratracheally administered with 2 mg/kg CIPS. After 3 days, mice were sacrificed and blood collected. For biochemical analysis, a 200 μl of whole blood was centrifuged to obtain plasma for 10 min at 3000 rpm and 4°C. Lung tissue was excised and fixed in 10% formalin for 24 h, and sectioned for hematoxylin and eosin (H&E) staining. All tests were performed by Wuhan Servicebio Technology Co.

### Supporting results and discussion

#### SARS-CoV-2 pseudoviruses and cell infection

The interaction of the SARS-CoV-2 Spike protein with the ACE2 protein on human cells is the key for SARS-CoV-2 cell entry^19,20^. To screen possible anti-SARS-CoV-2 agents without risk of infection, we have established and validated two pseudoviruses, expressing the Spike protein of SARS-CoV-2 (SC2-P) and SARS (SARS-P), using a replication-defective HIV lentivirus^21^. We constructed the SARS-CoV-2 and SARS Spike protein (S)-containing plasmids, and showed the expression of corresponding S protein after mammalian cells’ transfection (fig. S1a). This established S protein plasmids were used to generate psedoviruses. By using a lentivirus bearing the glycoprotein of the pantropic VSV (VSV-G) as control, the validation of the pseudoviruses was obtained by demonstrating the effective expression of S proteins on SC2-P or SARS-P by immunoblot analysis (fig. S1b).

The endocytic process of infection with pseudovirus through the ACE2 receptor was further demonstrated. SC2-P or SARS-P successfully entered Vero-E6 cells but not the ACE2 knocked-down Vero-E6, while the VSV-G pseudovirus equally entered both cell types, suggesting an ACE2-dependent endocytosis of SC2-P and SARS-P (fig. S1c). With luciferase carrying pseudoviruses, it was possible to show the pseudovirus entry in Vero-E6 cells (fig. S1d) and ACE2 stably transfected 293T cells (fig. S1e) but not in non-susceptible ACE2-negative 293T cells. Further immunofluorescent (IF) analysis also proved that the ACE2-overexpressing 293T cells were susceptible to infection by SC2-P and SARS-P (fig. S1f). The 3D image shows the ACE2 co-localization with SC2-P and SARS-P (fig. S1g-h), supporting the ACE2-dependent pseudoviral entry. The time course of pseudoviral infection showed a maximal cellular entry at 2 h (fig. S1i-l, fig. S2).

#### Screening of nanomaterials for potential inhibition of SARS-CoV-2 infection

Different NMs were screen for their potential capacity to inhibit cell infection by SARV-CoV-2. The characteristics of these NMs are presented in fig. S3a. Their potential inhibitory effects on SARS-CoV-2 infection were assessed with LUC gene-containing pseudovirus. As shown in fig. S3b, CIPS showed the strongest inhibitory activity on SARS-CoV-2 infection in the absence of any direct toxic effect. MoS_2_ presented limited anti-SARS-CoV-2 effect, while graphene oxide (GO) and Au-SC particles had variable concentration-dependent effects. Although black phosphorus (BP) showed some inhibitory effect on SARS-CoV-2 infection, this is most likely due to its significant cytotoxicity. Therefore, we chose CIPS for additional studies.

#### Evaluation of the contribution of ions in the antiviral effect of CIPS

We examined whether copper ions released from CIPS and related compounds could have anti-viral activity. Cu^+^, Cu^2+^ and In_2_S_3_ proved unable to exert antiviral activity on SC2-P infection in ACE2/293T cells (fig. S8a-d). We also used EDTA to chelate these ions in CIPS (fig. S8e). EDTA-treated CIPS retained its antiviral effect on SC2-P infection, suggesting that the antiviral capacity may does not depend on metal ions.

#### Molecular dynamics simulations of the interaction of RBD with nanomaterials

##### *Selection of CIPS surface for MD simulation*

We selected one surface of the CIPS NS to run MD simulation. Due to the heterogeneous distribution of Cu on the two sides of CIPS, surface charge on each side is different. The surface containing less Cu is more electronegative. According to the Adaptive Poisson-Boltzmann Solver (APBS) electrostatics calculations^22^, the surface of the RBD domain in the S protein is electropositive, thereby promoting adsorption on the negative CIPS side (fig. S13-S14).

Based on the surface potential of RBD calculated by APBS method, the conformation of RBD exhibits five sides (fig. S15a, b). The blue surface is the side with α1 to α3; the red surface is the side with α4, α5; the yellow surface is the side with β5, β6; the purple surface corresponding to the other right-angle side, and the green surface corresponding to the hypotenuse side. The total interaction energy between protein and CIPS at the final binding state was used to evaluate the adsorption priority onto each side of CIPS for 3 independent runs. The results show an average interaction energy of 1167.554±206.232 kJ/mol for RBD binding to the more electronegative CIPS surface, while the energy for RBD binding to the more positive CIPS surface is -761.604±185.060 kJ/mol. Thus, the electronegative surface containing 15% Cu was selected for the simulations (fig. S14).

##### *Selection of RBD side for MD simulation*

As mentioned above, RBD shows five sides on the surface, shown with five different colors (fig. S15a, b). The electrostatic potential maps of RBD correspond to the initial interaction of CIPS and RBD (fig. S15c). We preformed five independent runs for each initial simulation state (each of the five sides) of RBD approaching the electronegative CIPS surface. After 100 ns simulations, the 25 final states were collected for clustering. The clustering was based on RMSD and the cutoff was 8 Å. A typical configuration, the number of configurations in the cluster and the interaction energy of the typical configuration for each cluster are listed in Table S4. Although the results show that Cluster 2 has the highest interaction energies, this binding side of RBD in Cluster 2 is actually unexposed, as it is inside the S protein and thus the interaction of this side with CIPS is most likely not possible. Among other potent binding sides of RBD, the RBD-CIPS interaction energy in Cluster 1 was the lowest (fig. S15d, Table S4). This RBD binding side is the same involved in the RBD and interaction with ACE2. Thus, this configuration (Cluster 1) was selected for the RBD-CIPS binding simulation.

##### *Binding interface between CIPS and RBD*

To verify the contribution of amino acid residues to the RBD binding to CIPS, we have decomposed the interaction energy into different types of residues according to the amino acid side chains: amino (Arg and Lys); methyl (Ile, Leu, Ala and Val); benzene (Phe, Pro and Trp); hydroxyl (Ser, Tyr, and Thr); thiol (Cys); carboxyl (Asp and Glu); and Gly (Table S5). In most cases, the positively charged residues contributed to the majority of interaction energies.

The snapshots of a typical trajectory in Cluster 1 are reported in Fig. 5a and Movie S1. In the initial state, RBD is far from the CIPS layer. At 5.7 ns, the residues Asn 481 and Gln 498 take contact with CIPS. From 12.8 ns, the number of contacting residues increases dramatically, and the RBD begins to tilt. At 20.1 ns, 23 residues contact the CIPS surface. Afterwards, the RBD conformation remains stable. With the increase of contacting atoms in RBD, the interaction energy decreases and stays stable until 100 ns (Fig. 5a and Table S4). At the final state of the adsorption, the positively charged residues contribute to most interaction energies, and other polar residues ranked second. The hydrophobic residues also make large contribution to the interaction energies. The comparison on the interaction energies of different functional groups of RBD is reported in Fig. 5b.

Large conformational changes in RBD are observed after its adsorption onto the CIPS surface, especially in the binding region. The root-mean-square deviation (RMSD) of RBD was 3.69 Å. In the binding region (residue 445 to 456 and 473 to 501), the RMSD was 4.68 Å, while the RMSD of the other part was 2.54 Å. According to the crystal structure of the RBD-ACE2 complex (PDB ID: 6M17), there are 11 key residues that contact ACE2, among which six residues also contribute to the binding of RBD to CIPS (Table 1 in main text and Table S5). Among the 11 residues, six of them are located in the turn region, two of them in the β sheet region, and the others belong to coil region. Finally, the adsorption of CIPS to RBD not only changes the secondary structure of RBD but also blocks the binding of RBD to ACE2.

##### *Binding of CIPS with mutated N501Y RBD*

To check whether CIPS can still bind with mutant SARS-CoV-2, we performed 40 ns MD simulation to study the interaction of mutant N501Y RBD with CIPS. The potential energy between the SARS-CoV-2 RBD and ACE2 is -740 ± 70 kJ/mol^23^. When Asn (N501) residue is mutated to Thr (Y501), the local structure around N501 changes (fig. S16a-b) and the binding energy of Asn (N501) with CIPS significantly reduces from 49.71 kJ/mol to -70.68 kJ/mol of Tyr (Y501) (fig. S16c). The lost of Asn caused the decreased binding energy of Lys, Arg, Ser, Thr and Pro with CIPS, and the presence of Tyr increased the binding energy of Asp, Gly, Val and Phe (fig. S16b). Thus, the local mutation affects the binding energies of other residues with CIPS. The potential energy between the mutant N501Y RBD and CIPS changed to -1218.06 ± 37.75 kJ/mol (fig. S16d-e), which is the same as the energy of wild type RBD with CIPS, -1217.73 ± 49.97 kJ/mol at 40 ns (fig. S16d). This result suggests that CIPS still maintains high affinity with the mutant N501Y RBD. Thus, CIPS can bind to the mutant SARS-CoV-2 N501Y RBD that efficiently inhibit the infection of mutant SARS-CoV-2.

##### *Binding of nanosheets to RBD*

By comparing three types of 2D NS, both experiments (Fig. 4d and fig. S12) and theoretical simulations show that CIPS has the strongest affinity for the RDB domain (fig. S17). In the simulation, the interaction energy between CIPS and RDB can be attributed to electrostatic energy (Coulomb interaction) and vdW energy (Lenard Jones interaction) (Table S6). The electrostatic interaction is directly related to the atomic partial charge, while the vdW interaction intensity is directly related to the parameter ε, which determines the well depth of Lenard Jones potential. In the same protein adsorption mode, both of them are related to the surface atomic density of NS. Therefore, we first investigated the surface atomic density of CIPS and MoS_2_, which both have flat surfaces. The average surface atomic density of CIPS is 12.863/nm^2^ (in which S is 9.326/nm^2^ and P is 3.109/nm^2^) and the average surface charge density is 6.262/nm^2^ (in which S is 3.711/nm^2^ and P is 2.468/nm^2^). However, there are only S atoms on the MoS_2_ surface, and the atomic density of the surface is 11.725/nm^2^, the average surface charge density is 4.406/nm^2^. In addition to S, there are positively charged P atoms on the CIPS surface, which also interact with some negatively charged amino acids on RBD *via* electrostatic attraction. Thus, CIPS shows stronger electrostatic force in its interaction with RBD compared to MoS_2_ that only has negative charges. Compared with GO, O and H atoms are exposed on the surface, and the surface atomic density is 10.288/nm^2^, and the average surface charge density is 1.512/nm^2^. Therefore, the electrostatic attraction of CIPS for RBD is much stronger than that of GO and MoS_2_. As for the vdW energy, the well depth of Lenard Jones potential ε of S and P on CIPS are also larger than O and H on GO. In summary, the differences between CIPS and other NS in adsorption affinity (Experimental results, Fig. 4d and fig. S12) are consistent with the simulation results that show the distinct interaction energy, contact atom number, and contact area on NS, which are due to the types and distribution (*i.e.,* topological structure) of the elements in these NMs.

#### Effect of CIPS on SARS pseudovirus infection

To explore the specificity of the antiviral capacity of CIPS we have evaluated its effects on infection by the SARS pseudovirus (SARS-P). The antiviral effect of CIPS is evident also in this case (fig. S19), supporting the hypothesis that CIPS interferes with the S protein RBD binding to ACE2, a mechanism of entry shared between SARS and SARS-CoV-2.

#### CIPS biocompatibility and biosafety

CIPS biocompatibility and safety were assessed both *in vitro* (fig. S5) and *in vivo* (fig. S20). No cytotoxicity could be observed with various cell lines after CIPS treatment. *In vivo*, no adverse effect was detected 3 days after acute intratracheal administration of CIPS at 2 mg/kg CIPS, both systemically (blood parameters) and in the lung (histological examination).

### Supporting figures and tables

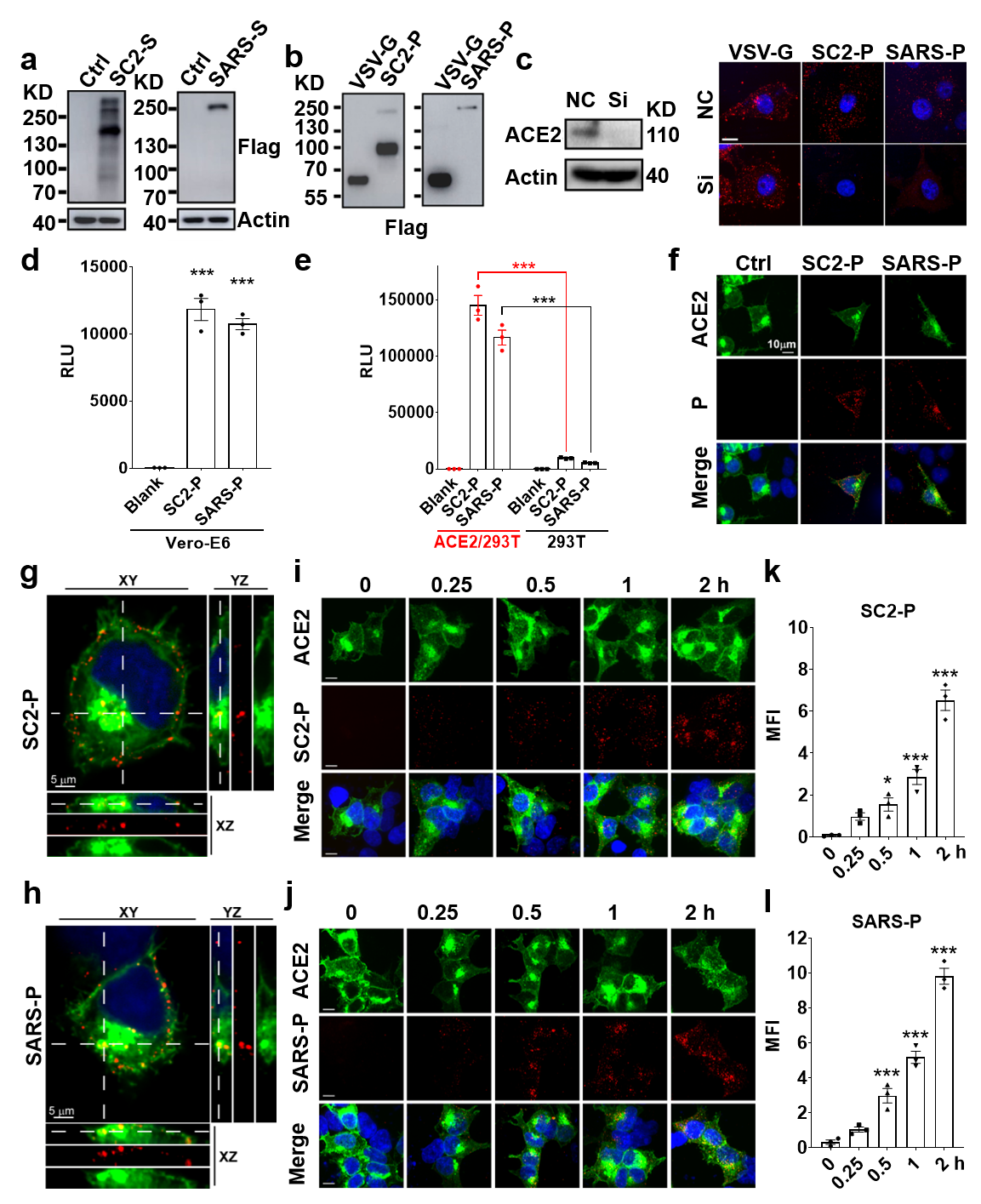

**Fig. S1. Establishment of** **pseudoviruses expressing the Spike protein of SARS-CoV-2 and SARS.** a-b) HEK-293T cells were transfected with empty vector (Ctrl) or SC2-S-Flag/SARS-S-Flag vectors. The S proteins of SARS-CoV-2 (SC2-S) and SARS (SARS-S) were detected by WB with anti-Flag antibody in cells lysates (a) and purified pseudoviruses (b). VSV-G pseudovirus was used as control. c) Infection of control and ACE2 knocked-down Vero-E6 cells with the SARS-CoV-2 pseudovirus (SC2-P) or with the SARS pseudovirus (SARS-P). NC: Vero-E6 cells transfected with scrambled siRNA (negative control); Si: ACE2 siRNA knocked-down Vero-E6 cells. ACE2 expression in control and ACE2 knocked-down cells was assessed by WB (left), and by immunofluorescent staining (right) of cells infected with SC2-P or SARS-P. VSV-G serves as negative control. d-e) Infective capacity of SC2-P and SARS-P on Vero-E6 (d) and 293T (e) cells, assessed by pseudovirus-dependent luciferase activity. Blank: mock control; SC2-P: LUC-expressing SARS-CoV-2 pseudovirus; SARS-P: LUC-expressing SARS pseudovirus. Data are from one experiment representative of three performed. Data are presented as mean ± SEM of technical triplicates. f) Co-localization of SC2-P/SARS-P with ACE2. ACE2-GFP overexpressing 293T cells (ACE2-OE) were infected with SC2-P or SARS-P for 2 h. Green: ACE2-GFP; Red: pseudoviruses labeled with anti-Flag antibody. g-h) 3D confocal images of ACE2-OE cells infected with SC2-P and SARS-P. i-j) Time course of ACE2-OE cell infection by SC2-P and SARS-P, assessed by immunofluorescent staining. k-l) Mean fluorescent intensity (MFI) quantitative analysis of acquired image (as shown in i and j). Data are representative of three independent experiments, and presented as mean ± SEM of three representative cells. *, *p*<0.05; ***, *p*<0.005 by ANOVA.

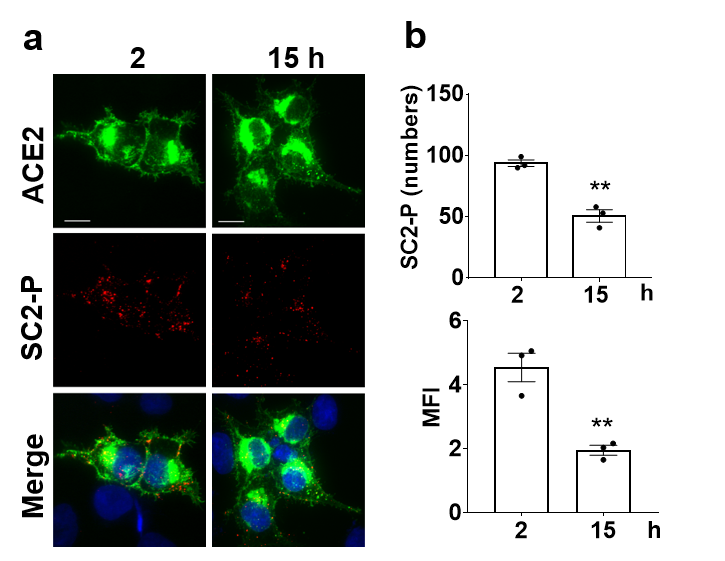

**Fig. S2. Infectivity of SC2-P at prolonged times.** a) Immunofluorescence images of infection of ACE2-OE cells exposed to SC2-P for 2 and 15 h. Green: ACE2; red, SC2-P. b) Quantitative evaluation of pseudovirus infectivity observed in panel a. The SC2-P particle number within cells was counted (cells ≥ 3) (top) and the mean fluorescent intensity (MFI) was assessed (bottom). Data in b are representative of three independent experiments, and are presented as mean ± SEM of three representative cells. **, p<0.01 by Student’s *t-*test.

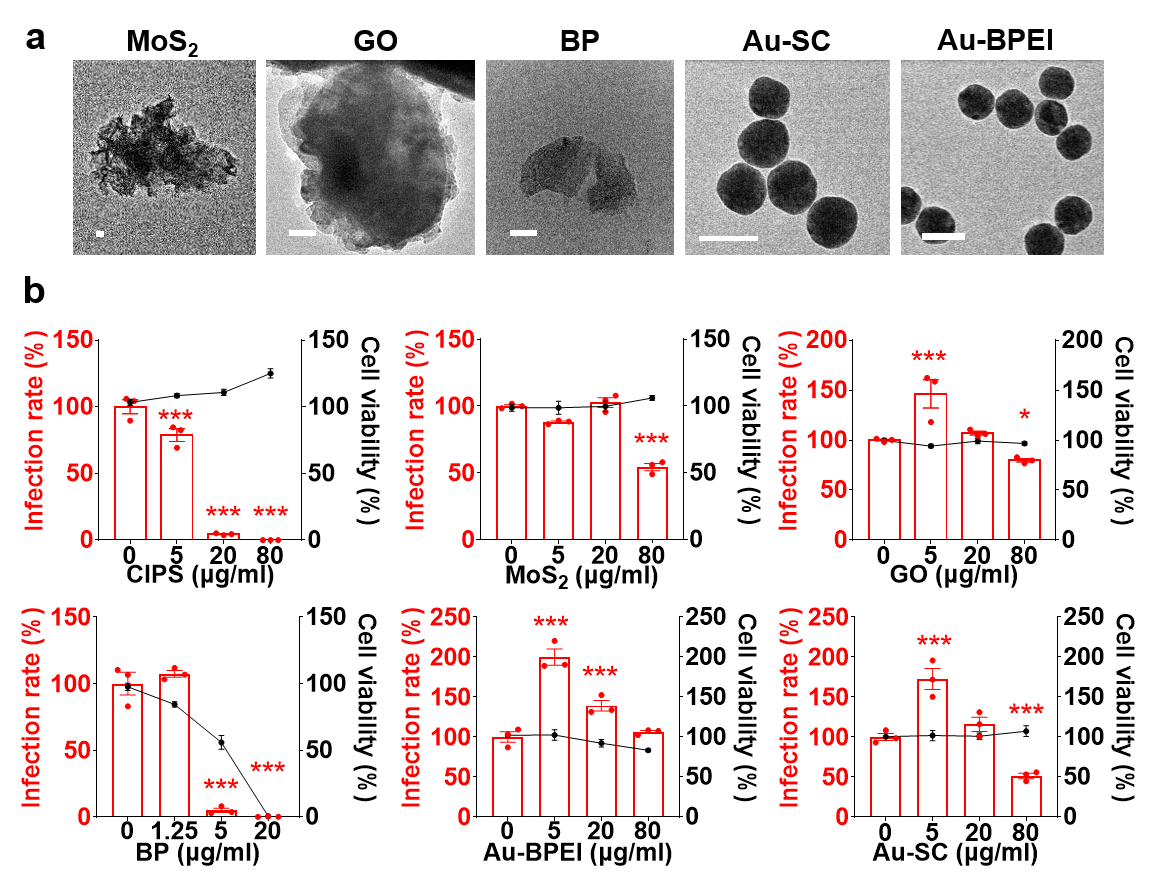

**Fig. S3. Screening of nanomaterials for potential inhibition of SARS-CoV-2 infection.** a) TEM images of the examined NMs. Scale bar = 50 nm. b) The effect of NMs on the infection rate (red) and cell viability (black) of ACE2/293T cells infected with SC2-P. LUC-containing SC2-P pre-incubated with different NMs was used for infecting ACE2/293T cells. Infection was evaluated on cell lysates as SC2-P dependent luciferase activity after 40 h, while the percentage of metabolically active viable cells was evaluated with the CCK8 assay after exposure to NM for 48 h. Data in b are presented as mean ± SEM of 3-5 technical replicates. *, *p*<0.05; ***, *p*<0.005 by ANOVA.

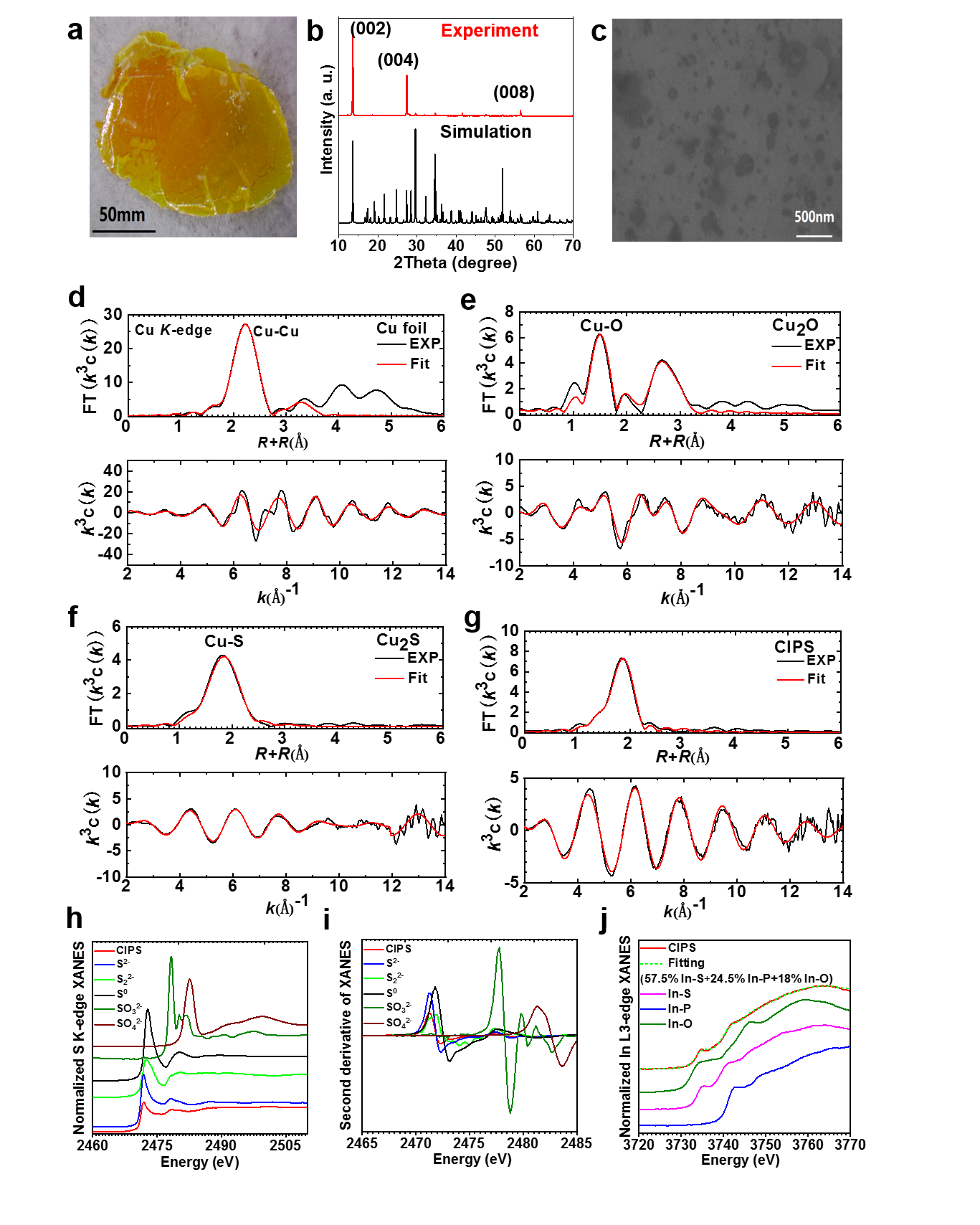

**Fig. S4.** **Characterization of CIPS.** a) Photo of bulk CIPS. b) CIPS crystal structures based on X-ray diffraction spectra for experimental and theoretical results. c) SEM image of exfoliated CIPS NS with a few layers. d-g) Coordination structure of Cu conf such as bond length and coordination number in reference samples (Cu foil, Cu_2_O), and CIPS NS based on experimental and fitting data for *κ*^2^ weighted EXAFS at the Cu *K*-edge for these chemicals. *ΔE* indicates the shift of the energy of *E*0. Fine coordination structure of Cu-Cu (d), Cu-O (e), Cu-S (f) in references and the bond with Cu atoms (g) in CIPS. h-j) Chemical forms of S and In within CIPS as characterized by X-ray near-edge structure (XANES). (h) Chemical form of sulfur in CIPS as measured by S K-edge XANES compared to reference samples including sulfide, disulfide, elemental sulfur, sulfite, and sulfate. (i) Derivative spectra of normalized XANES results for CIPS and the reference samples. (j) Chemical form of In in CIPS as measured by In L_3_-edge XANES and compared to reference samples including In_2_S_3_, InP and In_2_O_3_ and major components of chemical forms based on the linear combination fitting for CIPS XANES.

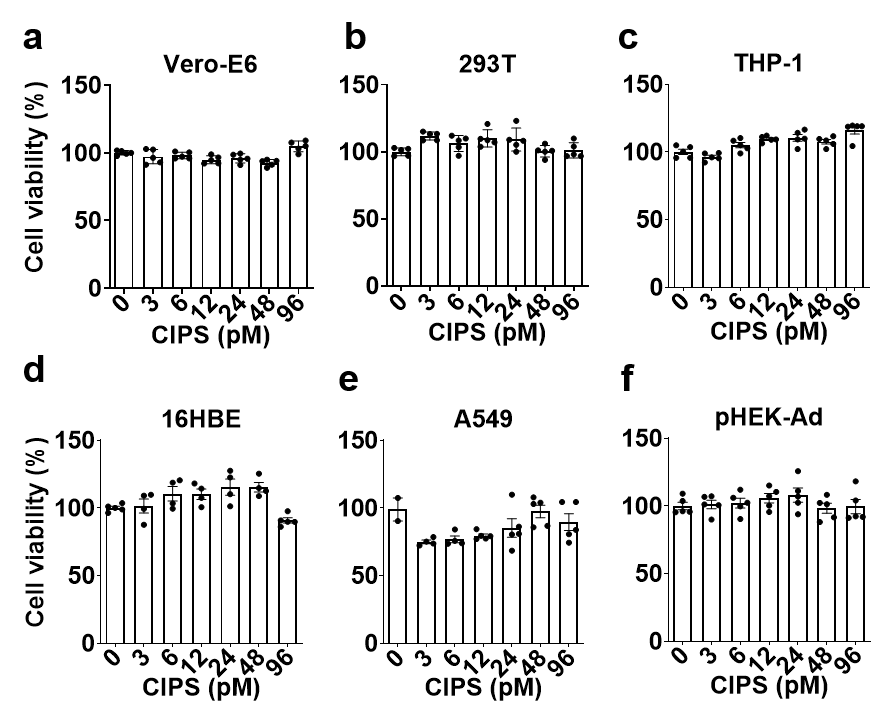

**Fig. S5. Cell viability after the exposure to CIPS.** Vero-E6, 293T, THP-1, 16HBE, A549, and pHEK-Ad cells were exposed to various concentration of CIPS for 24 h, and the number of viable metabolically active cells was evaluated with the CCK8 assay. Data are representative of two independent experiments, and present as mean ± SEM of four replicates. Results were analyzed by Student’s *t-*test with no significant difference.

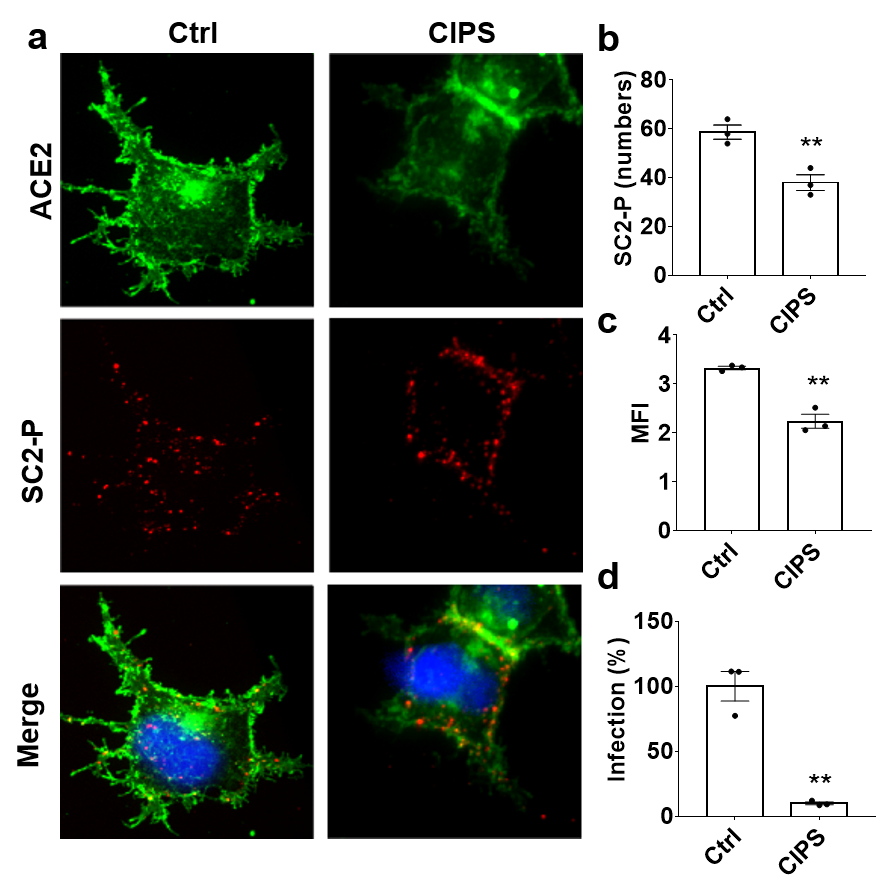

**Fig. S6. The inhibitory effect of CIPS on SC2-P infection.** a) ACE2-OE cells were incubated with SC2-P for 15 h in the presence or absence of 12 pM of CIPS. Green: ACE2-GFP; Red: SC2-Flag pseudovirus labeled with anti-Flag antibody. b-c) Quantitative evaluation of the effect of CIPS on SC2-P infectivity observed in IF (as in panel A). The SC2-P particle number within infected cells was counted (cells ≥ 3) (b) and the mean fluorescent intensity (MFI) assessed (b). d) SC2-P were pre-incubated with 12 pM CIPS for 2 h, and added to ACE2/293T cells for 15 h. After 40 h from infection, cells were lysed and SC2-P infection was assessed by pseudovirus-dependent luciferase activity. Data in b-d are representative of three independent experiments, and are presented as mean ± SEM of three representative cells. **, *p*<0.01 by Student’s *t-*test.

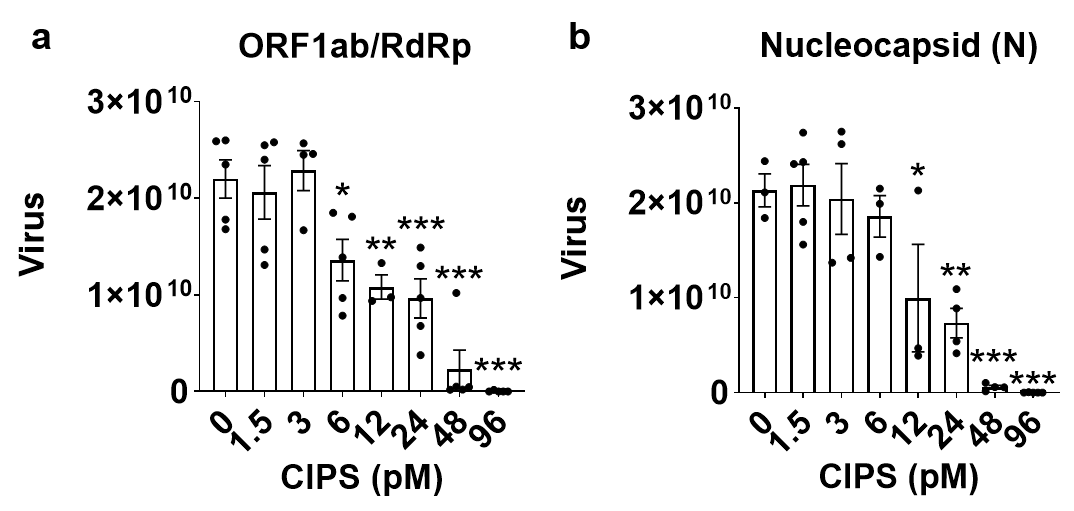

**Fig. S7. CIPS inhibits SARS-CoV-2 virus infection of Vero-E6 cells.** Vero-E6 cells were infected by SARS-CoV-2 virus for 1 h in the presence or absence of CIPS. Infective medium was then replaced with fresh medium containing CIPS and incubation prolonged for other 48 h. The virus in the 48 h-supernatant was quantitatively evaluated by real-time PCR for ORF1ab (a) and nucleocapsid (b). Data are representative of three independent experiments and are presented as mean ± SEM of 3-5 representative biological replicates. *, *p*<0.05; **, *p*<0.01; ***, *p*<0.005 by ANOVA.

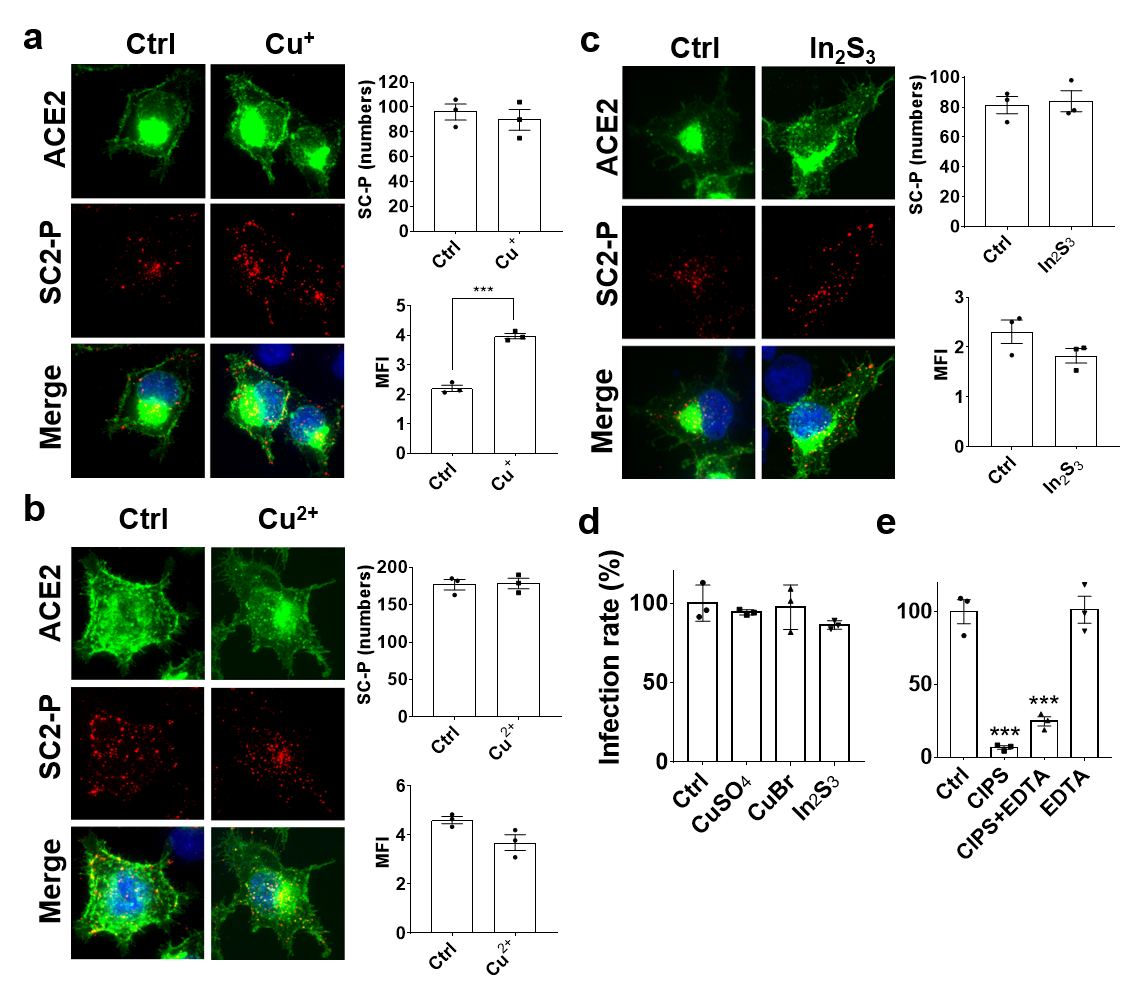

**Fig. S8. Effect of copper and indium-based compounds on the SC2-P infectivity.** a-c) Effect of Cu^+^ (a), Cu^2+^ (b), and In_2_S_3_ (c) on the SC2-P infectivity for ACE2-OE cells, assessed by IF. Quantitative evaluation was performed on acquired images for the number of SC2-P particles within cells (right top, cells ≥ 3) and for the mean fluorescent intensity (MFI, right bottom). SC2-P was exposed to CuBr, CuSO_4_ and In_2_S_3_ for 2 h, then added to ACE2-OE cells for 2 h. Green: ACE2-GFP; Red: SC2-Flag pseudovirus labeled with anti-Flag antibody. d) The effects of Cu^+^, Cu^2+^, In_2_S_3_ on SC2-P infectious activity in ACE2/293T cells assessed by luciferase activity. e) Effect of ion chelation on CIPS antiviral activity. CIPS (12 pM) was treated with/without EDTA, incubated with SC2-P for 2 h, then added to ACE2/293T cells for 2 h. SC2-P infection was assessed 40 h later as pseudovirus-dependent luciferase activity. Data in a-d are representative of three independent experiments and are presented as mean ± SEM of three representative cells. ***, *p*<0.005 by Student’s *t-*test.

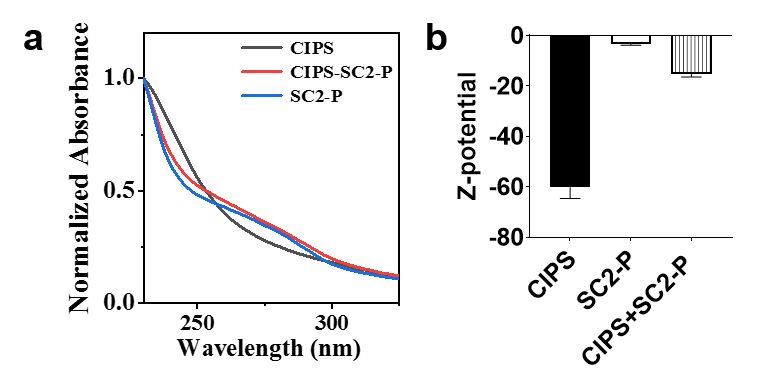

**Fig. S9. Characterization of the CIPS-SC2-P complex.** CIPS was incubated with SC2-P for 2 h and the CIPS-SC2-P complexes were isolated by centrifugation. **(**a) UV-VIS spectra and (b) Z-potential of CIPS, SC2-P and the SC2-P complexed with CIPS.

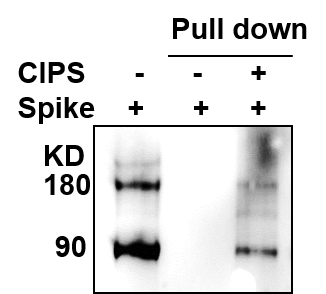

**Fig. S10. The binding** **of S protein with CIPS.** WB analysis indicating the binding of S protein with CIPS. The cell lysate of Spike-Flag protein expressed 293T cells was incubated with CIPS (12 pM) for 2 h, and further centrifuged to obtain the precipitates for WB analysis by anti-Flag.

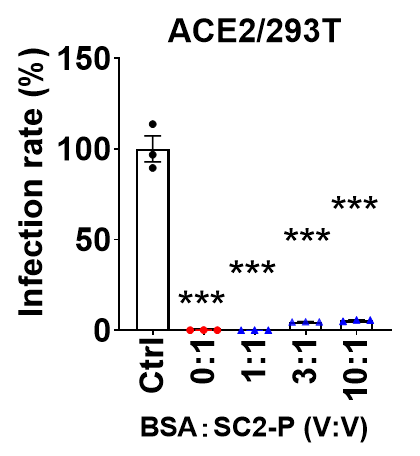

**Fig. S11. Antiviral activity of CIPS in the presence of BSA.** SC2-P was mixed with BSA and incubated with CIPS for 2 h, then used to infect ACE2/293T cells for 2 h. The infection by SC2-P was measured as intracellular luciferase activity after 40 h. Data are from one representative experiment of three performed, and presented as mean ± SEM of technical triplicates. ***, *p*<0.005 by ANOVA.

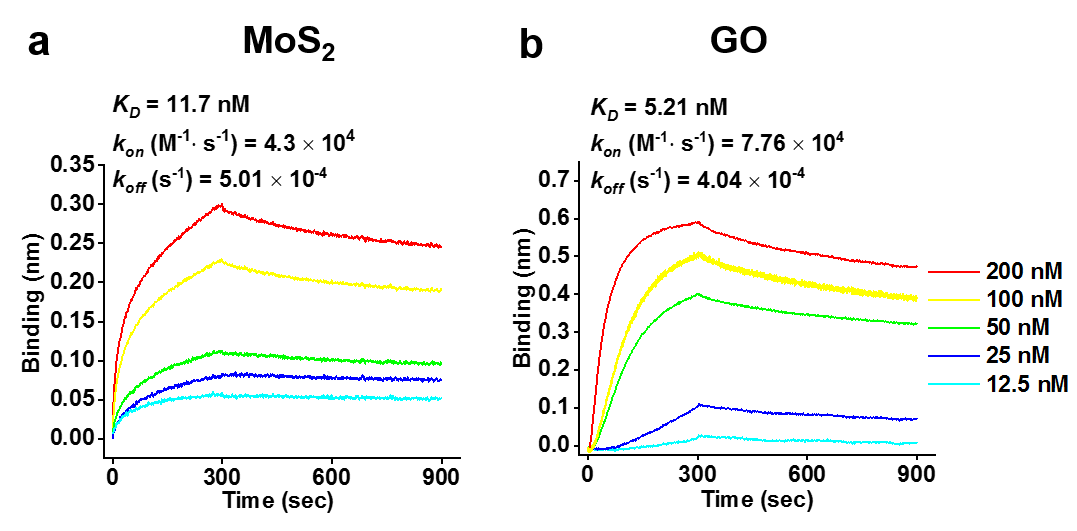

**Fig. S12. RBD binding affinity for graphene oxide (GO) and MoS2 nanosheets.** GO (20 μg/ml) and MoS_2_ (2 mg/ml) NSs were immobilized on the BLI sensor and immersed in RBD solutions at different concentrations.

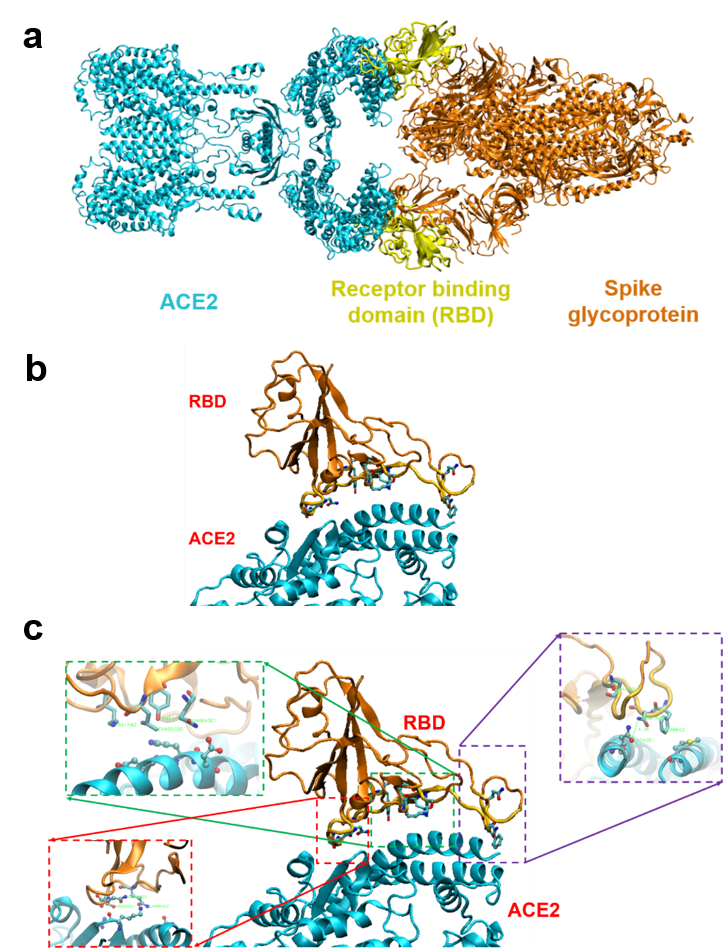

**Fig. S13. Structure of the RBD-ACE2 binding complex**. Structure of the binding interface between the RBD of the SARS-CoV-2 Spike protein and the human ACE2. (a) Crystal structure of a heterodimer of full-length human ACE2 (in cyan) and the S protein (in brown), in which the RBD is depicted in yellow. (b) Structure of the complex between the S protein RBD and ACE2. (c) Interface structure between RBD and ACE2. The insets show amino acid residues involved in the binding sites of RBD to ACE2. Amino acid residues and structure are listed in Table S4.

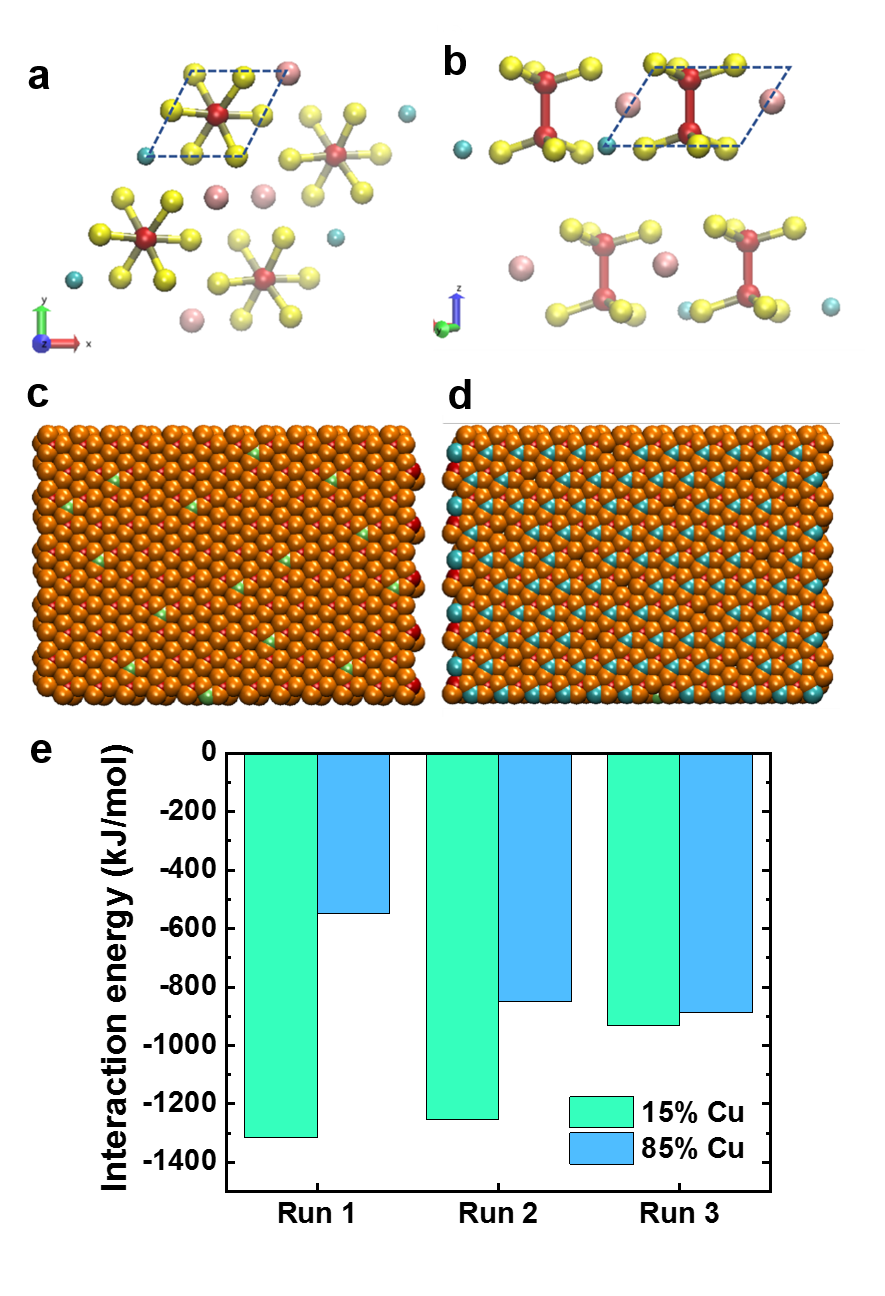

**Fig. S14.** **Structure of a CIPS lattice.** a-b) Structure of a CIPS lattice unit from a top view (a) and a side view (b). The parallelogram formed with the dash line represents a repeating unit containing one Cu atom, one In atom, two P atoms, and 6 S atoms. c-d) The two CIPS surfaces containing different number of Cu atoms. Left: electronegative surface containing 15% of the Cu atoms (green), used for studying the RBD-CIPS interaction (c). Right, opposite surface containing 85% of the Cu atoms (cyan) (d). e) The interaction energy between RBD and CIPS under three repeated simulation runs. Green columns represent the interaction energy between RBD and the CIPS surface with less Cu, while cyan columns represent the energy between RBD and the opposite, Cu-rich surface.

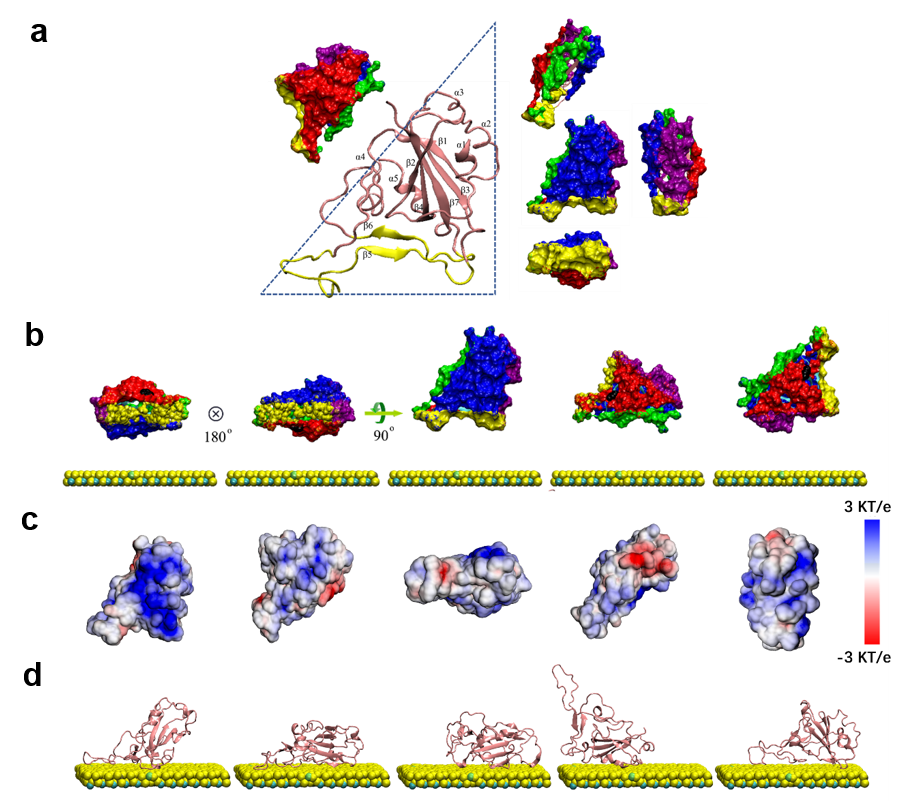

**Fig. S15.** **The conformation of RBD shown from different directions, and the five potential states at the start of the interaction with CIPS.** a) Conformation of RBD with its five sides depicted in different colors. The right angular dash line sketches the RBD contours. b) The initial state of RBD interaction with CIPS in MD simulations. Each one corresponds to one of five sides by which RBD approaches the CPIS plane. c) The electrostatic potential map of RBD corresponding to the initial states of the five RBD orientations, determined with APBS electrostatics calculations. The surface facing the reader corresponds to the side of the initial conformation facing CIPS. The blue and red colors represent the surface with positive and negative charges, respectively. d) Typical configuration for the binding of RBD to CIPS in five clusters. Configurations of RBD for the five clusters are based on 25 trajectories at 100 ns MD simulation.

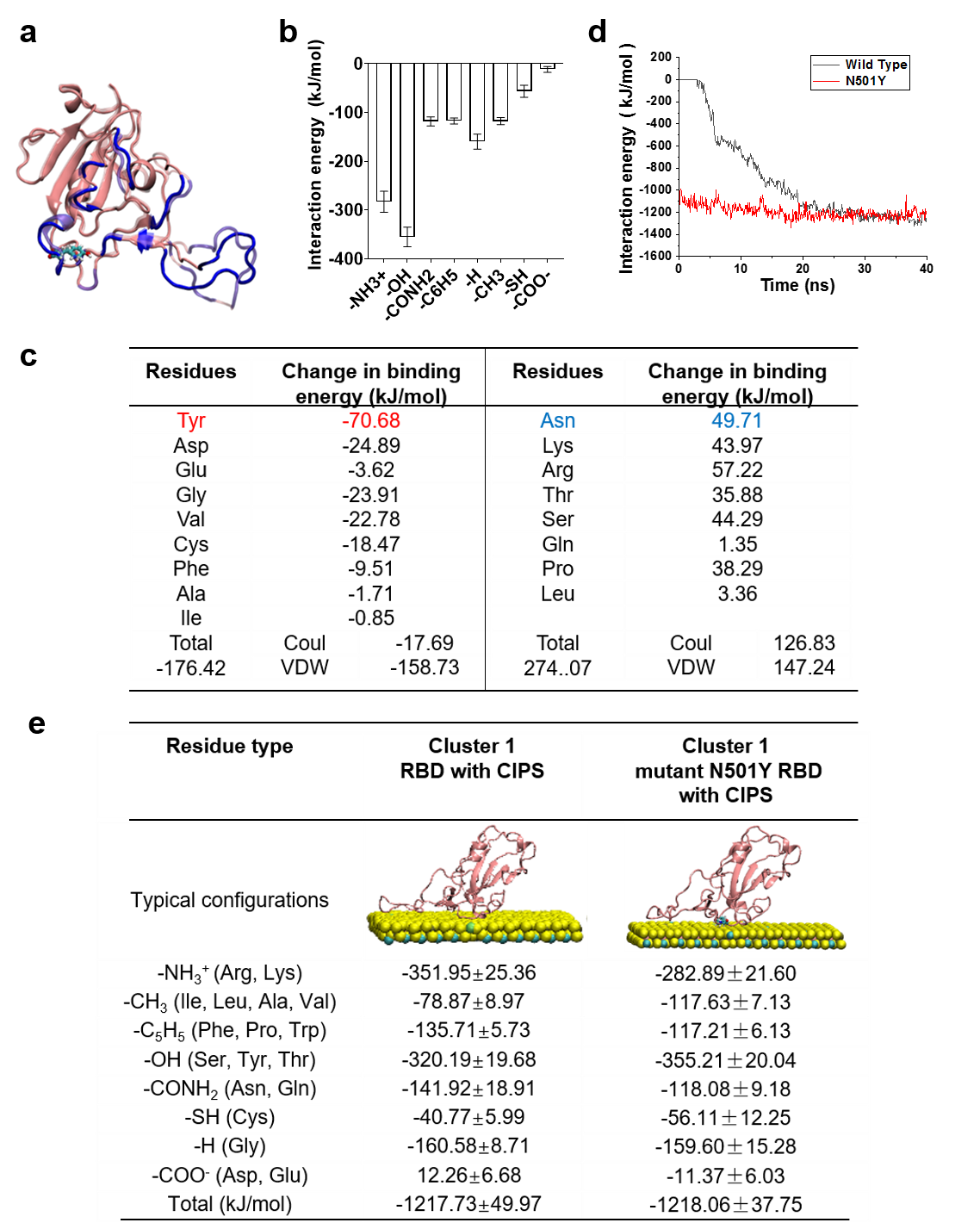

**Fig. S16.** **The interaction between mutant N501Y RBD and CIPS.** a) Representative configuration of mutant N501Y RBD and the adsorption interface. The cyan color indicates the mutant tyrosine residue that replaces asparagine (N501) within the RBD. The blue is the residues at the binding interface. b) The interaction energies of CIPS with the residue functional groups in the mutant RBD. c) The interaction energy changes (kJ/mol) between each RBD residue and CIPS before and after the N501Y RBD mutation. The major interactive forces include Coulomb interaction (Coul) and van der Waals (vdW) forces. d) The interaction energy of CIPS with wild type or N501Y RBD. e) Typical interaction configurations of CIPS with wide type RBD or mutant N501Y RBD. At 40 ns, the interaction energies and the amino acid residues mainly contributing to the binding force are shown for mutant RBD with CIPS. Before and after N501Y RBD mutation, the binding sites of amino acid residues are the same except the mutated one.

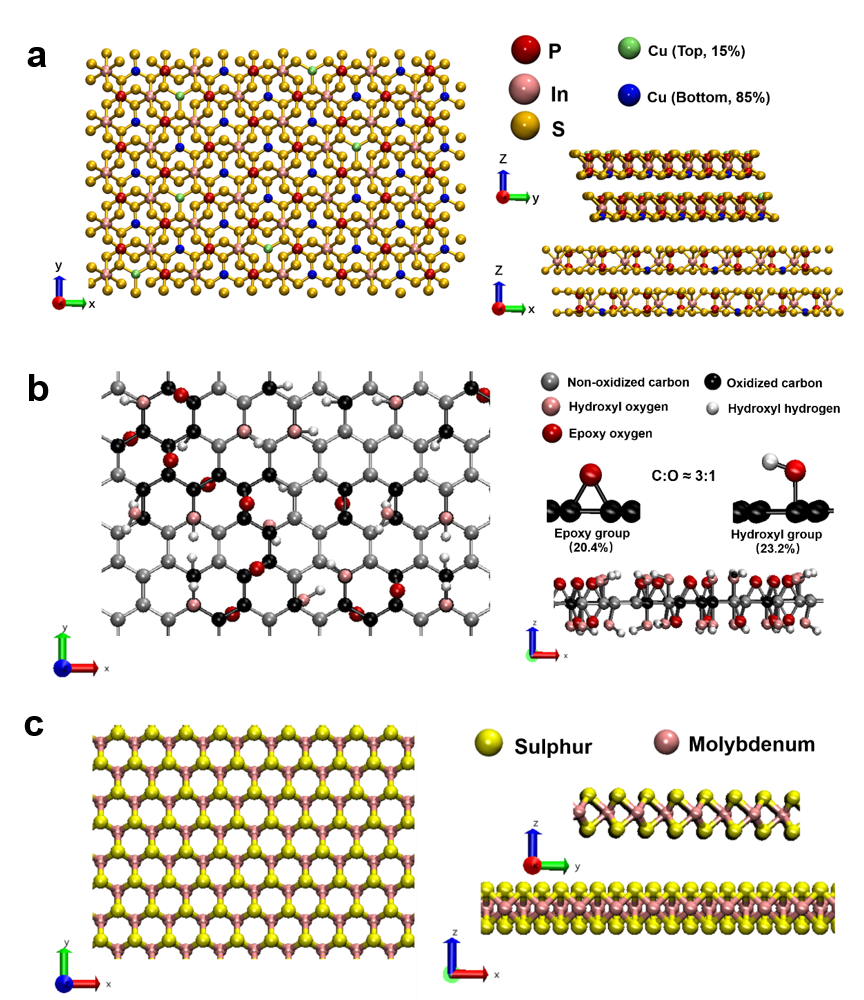
**Fig. S17.** **Configuration structure of CIPS, GO and MoS_2_ NS.** a) Surface structure of CIPS NS with a top and a side view. Phosphorus, indium, and sulfur atoms are shown in red, pink and yellow, respectively. b) Surface structure of GO with a top and a side view. Major functional groups of GO are distributed on the NS surface and are depicted in different colors. The ratio of -OH to =O was 2:1 and the GO sheets have a C/O ratio of 3:1. c) Surface structure of MoS_2_ NS with a top and a side view. Sulfur and molybdenum atoms are shown in yellow and pink, respectively.

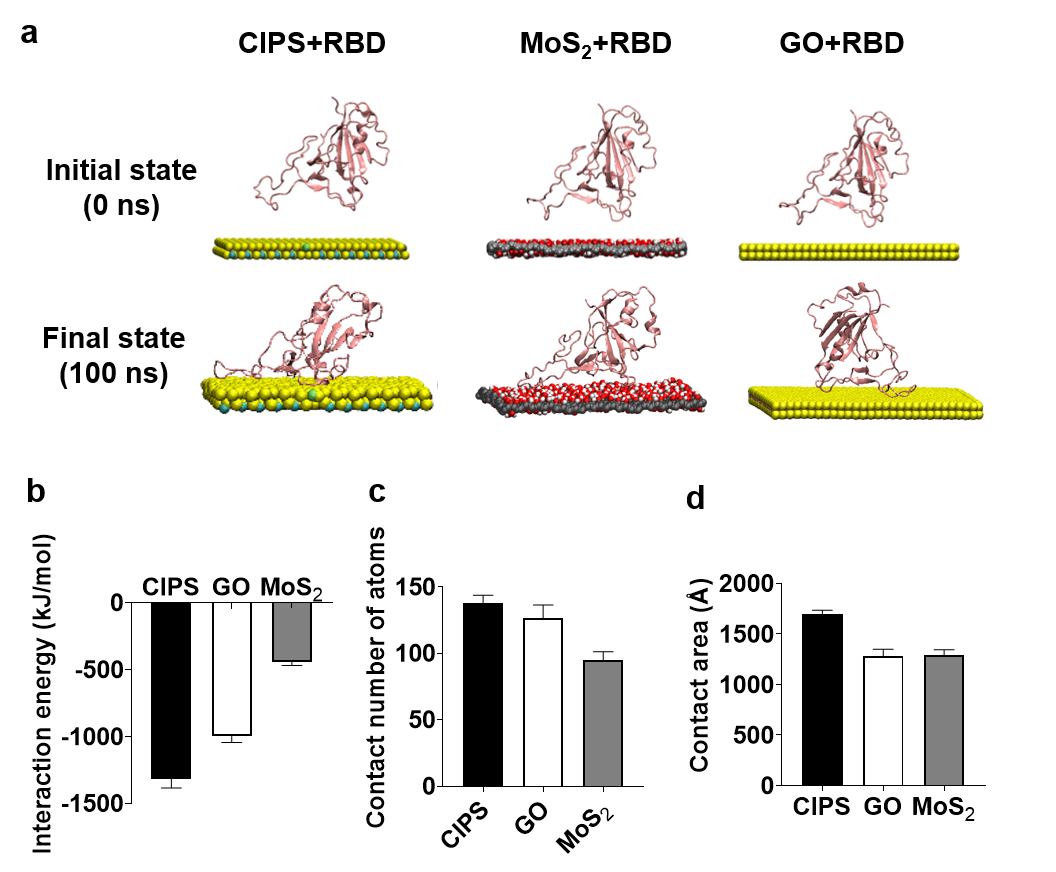

**Fig. S18. MD simulation of the binding interface between three types of nanosheets and RBD and the parameters for NS-RBD interaction.** a) The initial and the final states for the interaction of RBD with NSs based on 100 ns MD simulation. b) Interaction energy for RBD-NS binding at 100 ns from a representative trajectory. c) Number of RBD atoms contacting the NS at 100 ns. d) Contact area of RBD with the NS surface at 100 ns.

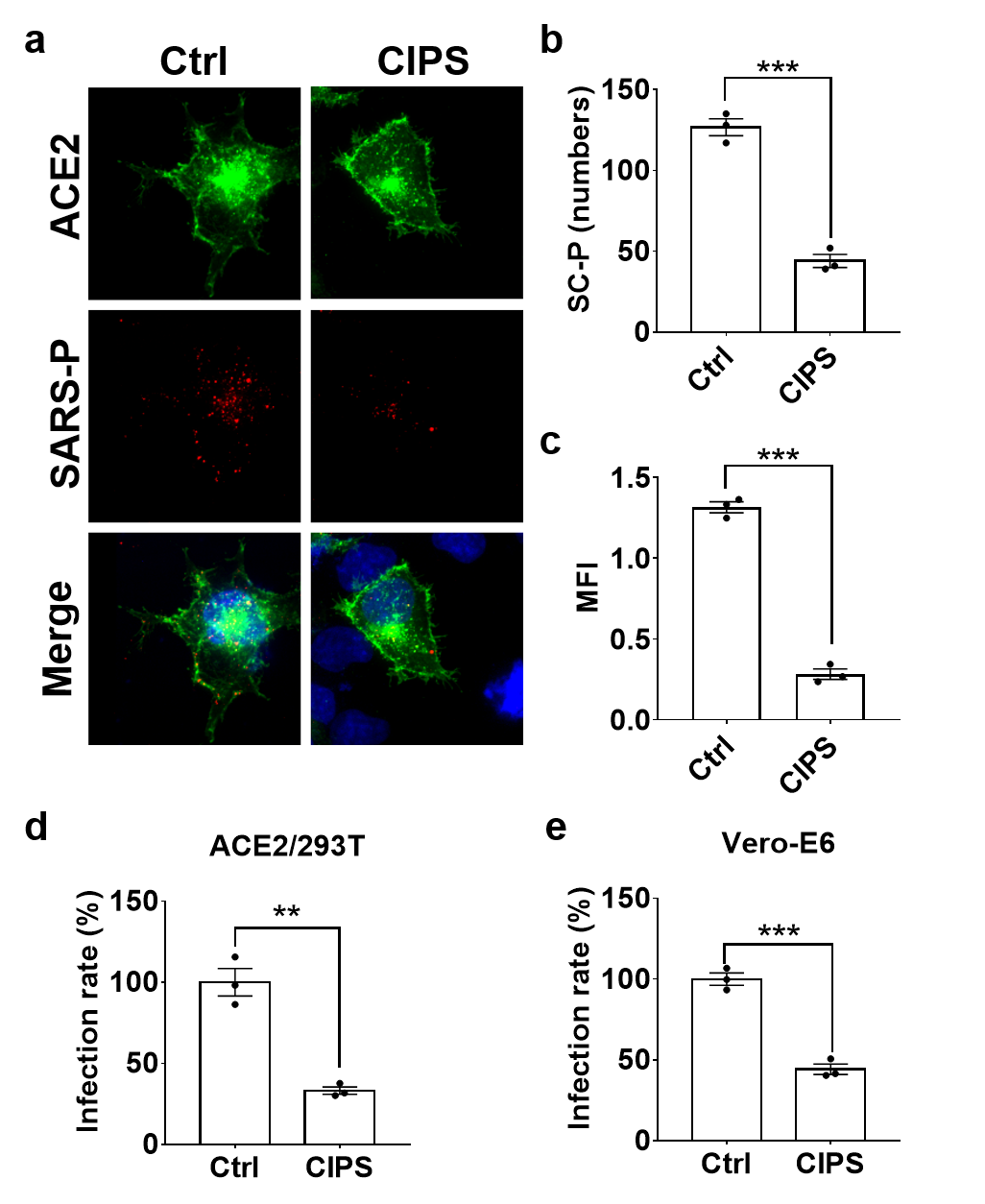

**Fig. S19. Antiviral activity of CIPS for SARS pseudovirus.** a) SARS-P pre-incubated with 12 pM CIPS for 2 h was added to ACE2-OE cells for 2 h. Green: ACE2-GFP; Red: SARS-Flag pseudovirus labeled with anti-Flag antibody. b-c) Quantitative evaluation was performed on acquired images in panel A for the number of SARS-P particles within cells (cells ≥ 3) (b) and for the mean fluorescent intensity (MFI) (c). Data are representative of three independent experiments, and presented as mean ± SEM of three representative cells. ***, p<0.005 by Student’s *t-*test. d-e) The inhibitory effect of CIPS on SARS-P infectivity for ACE2/293T (d) and Vero-E6 cells (e). SARS-P pre-incubated with 12 pM CIPS for 2 h was added to Vero-E6 or ACE2/293T cells for 2 h. SARS-P infection was assessed by pseudovirus-dependent luciferase activity 40 h later. Data are representative of three independent experiments, and presented as mean ± SEM of technical triplicates. **, *p*<0.01; ***, *p*<0.005 by Student’s *t-*test.

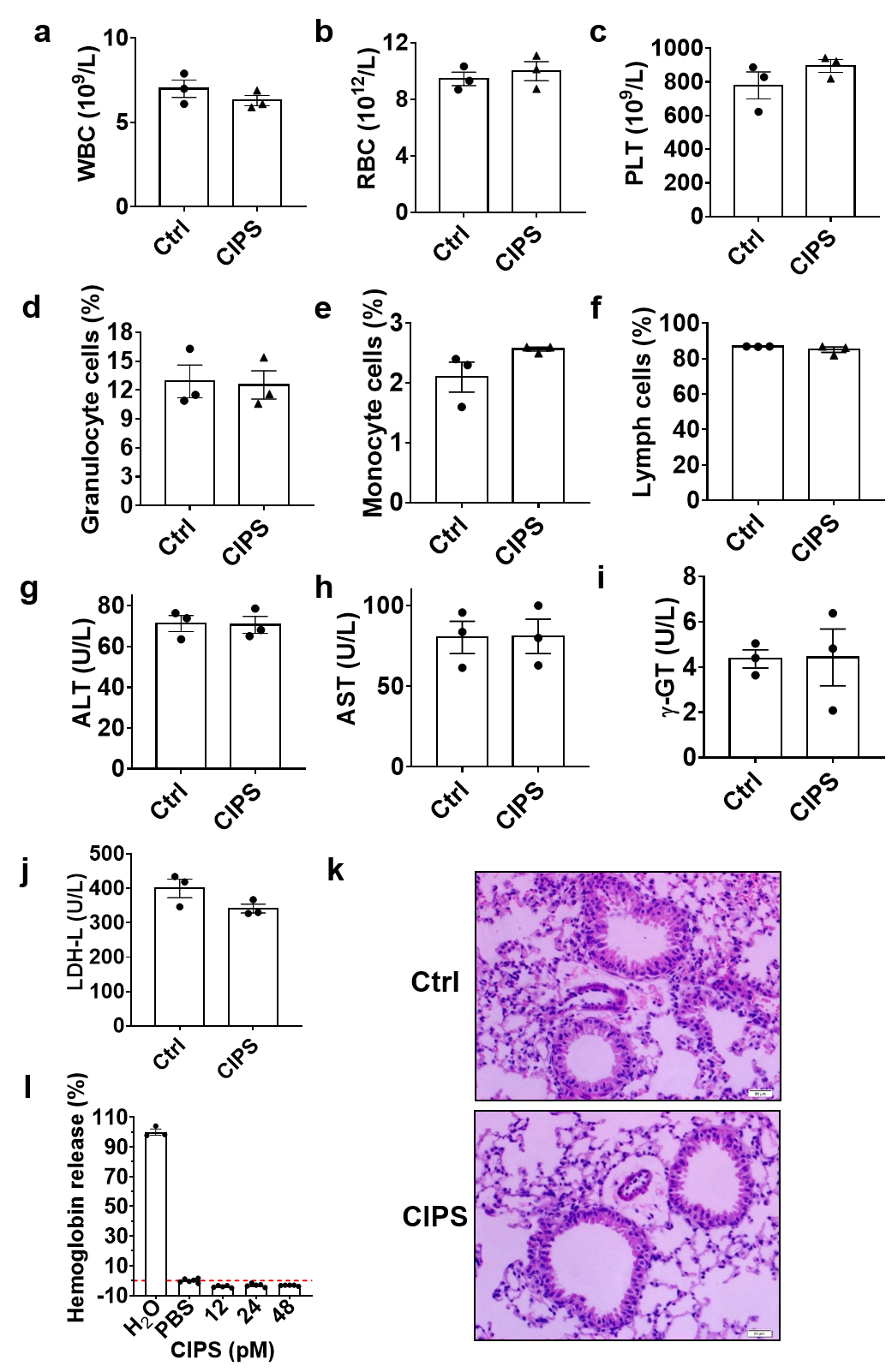

**Fig. S20. Evaluation of CIPS toxicity *in vivo*.** C57BL/6J mice were administered with CIPS intratracheally, and parameters assessed after 3 days. a-f) Blood cell counts. g-j) Serum chemistry panel. Data are presented as mean ± SEM of five mice. k) Hematoxylin and eosin (H&E) staining of representative lung sections. l) Hemoglobin release rate induced by CIPS. Data are presented as mean ± SEM of five representative replicates.

**Table S1.** The fine structure parameters fitted from *κ*^2^ weighted EXAFS at the Cu *K*-edge data for Cu foil, Cu_2_O, Cu_2_S, and CIPS NS

|  | Bond | *N* | *R* (Å) | σ^2^ (10^-3^Å^2^) | *ΔE* (eV) | *R*-factor |
| --- | --- | --- | --- | --- | --- | --- |
| **Cu foil** | 1) Cu-Cu1 | 12 | 2.544±0.005 | 9.32 | 1.89 | 0.02 |
|  | 2) Cu-Cu2 | 6 | 3.570±0.02 | 12.15 |  |  |
| **Cu_2_O** | 1) Cu-O | 1 | 1.854±0.009 | 0.2 | 9.67 | 0.02 |
|  | 2) Cu-Cu | 12.5 | 3.029±0.019 | 23.5 |  |  |
| **Cu_2_S** | 1) Cu-S | 2 | 2.273±0.024 | 10.75 | 2.76 | 0.02 |
|  | 2) Cu-Cu1 | 2 | 2.627±0.031 | 16.7 |  |  |
| **CIPS** | 1) Cu-S | 2.4 | 2.28±0.008 | 6.81 | 5.97 | 0.01 |
|  | 2) Cu-O | 3.2 | 2.24±0.048 | 66.5 |  |  |
|  | 3) Cu-Cu | 9.6 | 3.30 | 46.0 |  |  |

*N* indicates the coordination number. *R* indicates the bond length. *σ^2^* indicates the Debye-Waller factor. *ΔE* indicates the shift of the energy of *E_0_*.

**Table S2.** Chemical form of Cu in CIPS before and after exposure to THP-1 differentiated macrophages as determined by K-edge XANES.

| Samples | Chemical forms of copper | | | | | | | |
| --- | --- | --- | --- | --- | --- | --- | --- | --- |
|  | Cu (I) | | | Cu (II) | | | | |
|  | Cu_2_S | Cu_2_O | Cu-OOC- | CuS | | CuO | Cu-OOC- | |
| CIPS | 83% | 1.1% | 5.5% | 3.8% | | 0 | 6.5% | |
| 12 h uptake | 56.9% | 0 | 13.1% | 17.3% | 8.4% | | | 4.3% |
| 12 h degradation | 46% | 6.6% | 1.4% | 11.8% | | 12% | 22.2% | |
| 24 h degradation | 36.2% | 0 | 1.9% | 0.3% | | 18.2% | 43.4% | |
| 48 h degradation | 6.2% | 0 | 6.7% | 0 | | 17.7% | 70.7% | |

Macrophages were treated with 12 pM CIPS for 12 h and then cultured in CIPS-free medium to study the chemical forms of upon intracellular CIPS degradation.

**Table S3.** Binding affinity between CIPS and various proteins in the blood or in FBS, as determined by BLI.

| Sample | Binding affinity parameters | | |
| --- | --- | --- | --- |
|  | ***k_on_*** (M^-1 .^ s^-1^) | ***k_off_*** (s^-1^) | ***K_D_*** (nM) |
| CIPS + RBD | 1.73×10^5^ | 1.0×10^-7^ | <0.001 |
| CIPS + FBS | 1.61×10^4^ | 1.11×10^-4^ | 6.91 |
| CIPS + HSA | 6.29×10^4^ | 3.60×10^-4^ | 5.73 |
| CIPS + HDL | 2.33×10^5^ | 1.10×10^-4^ | 0.472 |
| CIPS + Fg | 1.65×10^5^ | 6.85×10^-5^ | 0.416 |
| CIPS + Tf | 6.56×10^4^ | 1.39×10^-4^ | 2.12 |
| CIPS + IgG | 1.78×10^5^ | 1.56×10^-4^ | 0.878 |

| Residue type | Cluster 1 | Cluster 2 | Cluster 3 | Cluster 4 | Cluster 5 |
| --- | --- | --- | --- | --- | --- |
| Typical configurations | 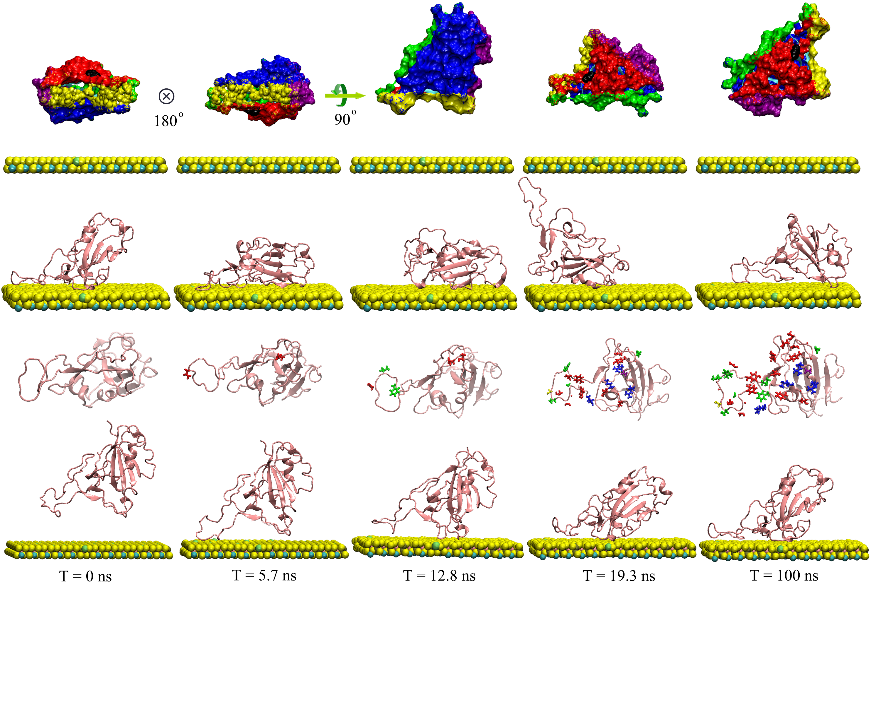 | 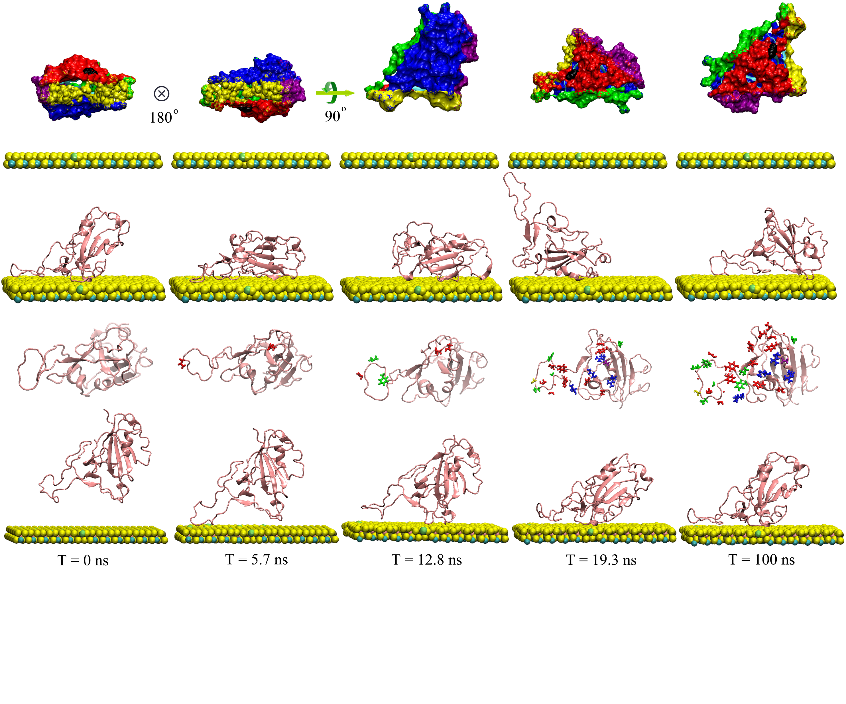 | 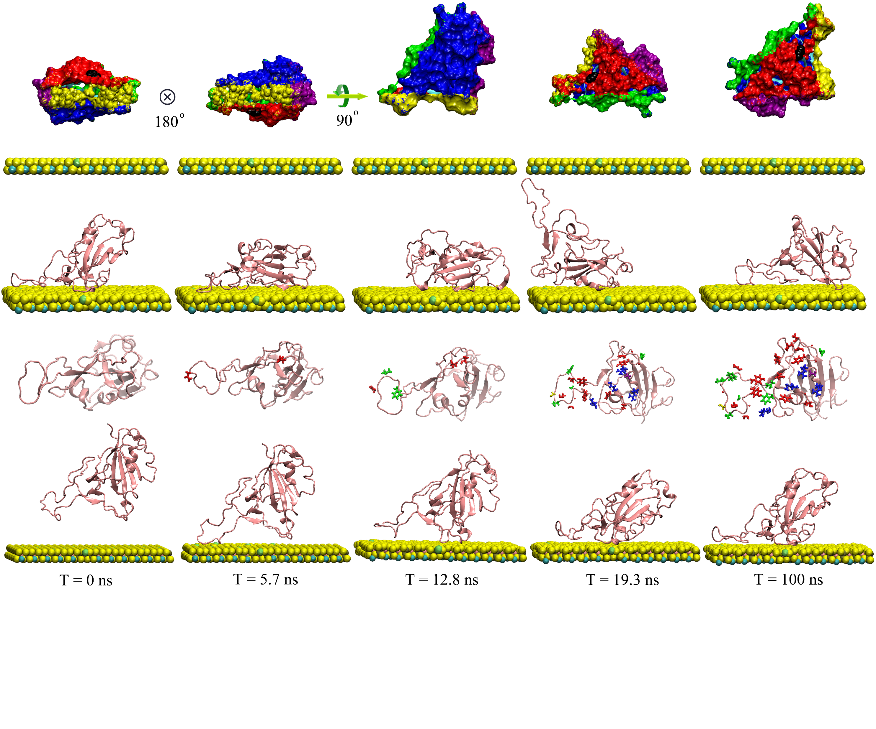 | 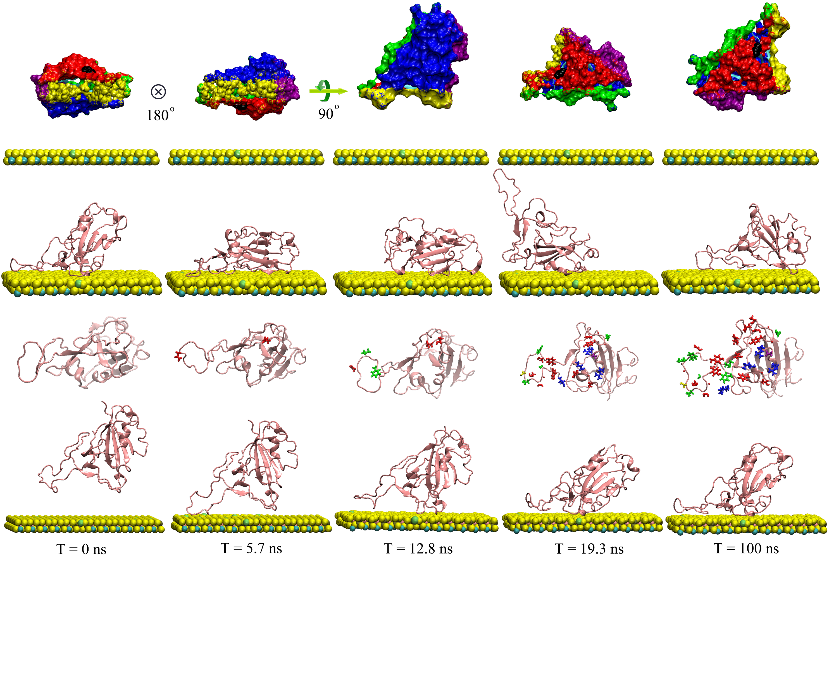 | 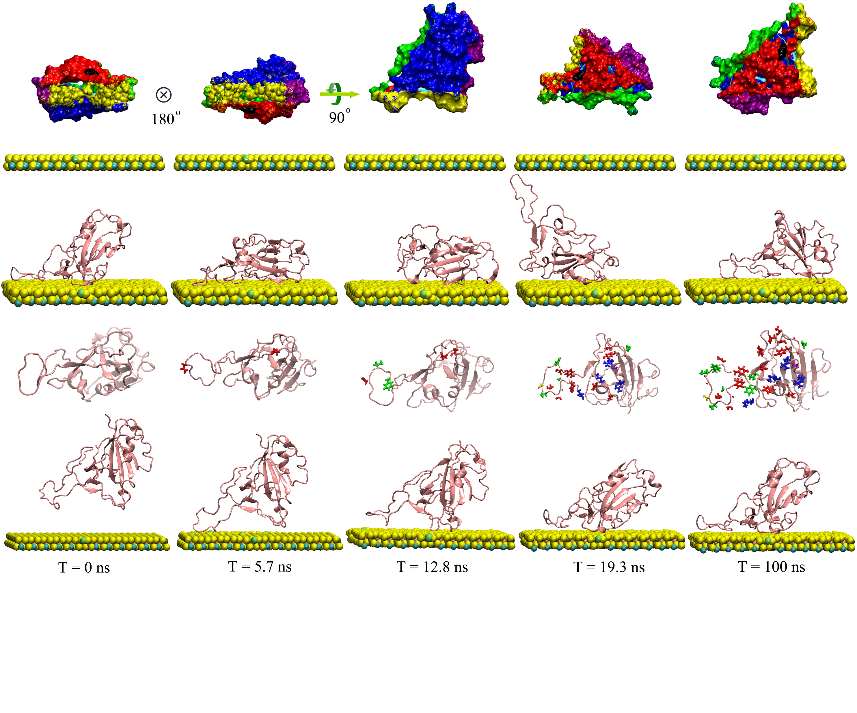 |
| -NH3^+^ (Arg, Lys) | **-384.09**±**19.38** | **-416.54**±**17.56** | -253.47±9.81 | -0.62±0.43 | **177.27**±**10.99** |
| -CH3 (Ile, Leu, Ala, Val) | -95.57±11.76 | -215.38±12.67 | -94.85±4.46 | -107.14±5.44 | -105.89±7.08 |
| -C5H5 (Phe, Pro, Trp) | -146.00±7.52 | -116.97±5.32 | -60.54±4.17 | **-176.73±11.78** | -92.89±7.14 |
| -OH (Ser, Tyr, Thr) | -364.72±24.87 | -181.85±10.54 | **-272.95±12.32** | -119.38±8.12 | -87.68±6.78 |
| -CONH2 (Asn, Gln) | -169.14±21.49 | -265.99±13.99 | -98.74±7.99 | -154.68±11.57 | -103.83±14.93 |
| -SH (Cys) | -37.60±14.63 | -115.48±7.71 | -6.37±0.87 | -45.74±7.14 | -55.44±7.64 |
| -H (Gly) | -135.64±12.51 | -109.11±10.07 | -49.00±7.72 | -6.58±0.96 | -44.92±4.48 |
| -COOH^-^ (Asp, Glu) | 17.14±5.04 | -57.15±8.22 | -4.54±0.63 | -5.59±2.72 | -10.21±6.13 |
| Total | -1315.62±66.85 | -1478.51±32.15 | -840.48±24.51 | -616.49±22.44 | -678.16±24.17 |

**Table S4.** Typical interaction configurations between RBD and CIPS in five clusters. The interaction energies and the amino acid residues mainly contributing to the binding force are shown.

**Table S5.** RBD amino acid residues contacting CIPS during the interaction of RBD with CIPS based on 100 ns MD simulations.

| **Amino acid type** | **T= 5.7 ns** | **T= 12.8 ns** | **T= 19.3 ns** | **T= 100 ns** |
| --- | --- | --- | --- | --- |
| Polar | ASN481 | ASN481 | ASN487, 501 | ASN481,487,**501*** |
|  | GLN498 | GLN498 |  | GLN493, **498*** |
|  |  | THR500 | THR415, 478 | THR415, 478, **500*** |
|  |  |  | TYR421, 473, 489, 505 | **TYR449***, 473, 489, 505 |
|  |  |  | GLY416, 476, 502, 504 | GLY416, 476, 482, 502, 504 |
|  |  |  | SER477 | SER477 |
|  |  |  | CYS480 | CYS480 |
| Charged |  |  | ARG403,408 | ARG403,408 |
|  |  |  | LYS417,458 | **LYS417***,458 |
|  |  |  | ASP405 | ASP405 |
| Hydrophobic |  | PHE486 | PHE486 | PHE456, **486*** |
|  |  | VAL483 | VAL503 | VAL483, 503 |
|  |  |  | PRO479 | PRO479 |
|  |  |  | ALA475 | ALA475, LEU455 |

* These residues that adsorbed by CIPS also interact with ACE2.

**Table S6.** Partial charges, σ and ε for atoms used in the MD simulation to study the molecular adsorption of RBD onto three types of NS.

| NS | Element | Charge | σ（nm） | ε（kJ/mol） | Reference |
| --- | --- | --- | --- | --- | --- |
| CIPS | Cu | 0.212 | 0.3495 | 0.0209 | ^7^ |
|  | In | 0.588 | 0.4463 | 2.5074 |  |
|  | P | 0.794 | 0.4147 | 1.2767 |  |
|  | S | -0.398 | 0.4035 | 1.1469 |  |
| GO | C (in Hydroxyl) | 0.180 | 0.340 | 0.3598 | ^24^ |
|  | O (in Hydroxyl) | -0.570 | 0.307 | 0.880 |  |
|  | H (in Hydroxyl) | 0.390 | 0.000 | 0.000 |  |
|  | C (in Epoxy) | 0.180 | 0.340 | 0.3598 |  |
|  | O (in Epoxy) | -0.360 | 0.300 | 0.711 |  |
| MoS_2_ | Mo | 0.760 | 0.255 | 0.543 | ^25^ |
|  | S | -0.380 | 0.350 | 1.1045 |  |

The ratio of -OH to =O was 2:1, and the GO sheets have a C/O ratio of 3:1. σ: the depth of the potential well (usually referred to as 'dispersion energy'); ε: distance at which the particle-particle potential energy V is zero (also referred to as the size of the particles).
